# The Dialysis Procedure Triggers Autonomic Imbalance and Cardiac Arrhythmias: Insights from Continuous 14-day ECG Monitoring

**DOI:** 10.1101/601542

**Authors:** Nichole M. Rogovoy, Stacey J. Howell, Tiffany L. Lee, Christopher Hamilton, Erick A. Perez-Alday, Muammar M. Kabir, Yin Li-Pershing, Yanwei Zhang, Esther D. Kim, Jessica Fitzpatrick, Jose M. Monroy-Trujillo, Michelle M. Estrella, Stephen M. Sozio, Bernard G. Jaar, Rulan S. Parekh, Larisa G. Tereshchenko

## Abstract

**Background:** In end-stage kidney disease the dialytic cycle relates to the rate of sudden cardiac death. We hypothesized that circadian, dialytic cycles, paroxysmal arrhythmias, and cardiovascular risk factors are associated with periodic changes in heart rate and heart rate variability (HRV) in incident dialysis patients.

**Methods:** We conducted a prospective ancillary study of the Predictors of Arrhythmic and Cardiovascular Risk in End Stage Renal Disease cohort (n=28; age 54±13 y; 57% men; 96% black; 33% with a history of structural heart disease; left ventricular ejection fraction 70±9%). Continuous ECG monitoring was performed using an ECG patch (Zio Patch, iRhythm) and short-term HRV was measured for three minutes every hour. HRV was measured by root mean square of the successive normal-to-normal intervals (rMSSD), high and low frequency power, Poincaré plot, and sample and Renyi entropy.

**Results:** Arrhythmias were detected in 46% (n=13). Non-sustained ventricular tachycardia (VT) was more frequent during dialysis or within 6 hours post-dialysis, as compared to pre-or between-dialysis (63% vs. 37%, P=0.015), whereas supraventricular tachycardia was more frequent pre-/ between-dialysis, as compared to during-/ post-dialysis (84% vs. 16%, P=0.015). In adjusted for cardiovascular disease and its risk factors autoregressive conditional heteroscedasticity panel (ARCH) model, VT events were associated with increased heart rate by 11.2 (95%CI 10.1-12.3) bpm (P<0.0001). During regular dialytic cycle, rMSSD demonstrated significant circadian pattern (Mesor 10.6(0.9-11.2) ms; Amplitude 1.5(1.0-3.1) ms; Peak at 02:01(20:22-03:16) am; P<0.0001), which was abolished on a second day interdialytic extension (adjusted ARCH trend for rMSSD −1.41(−1.67 to −1.15) ms per 24h; P<0.0001).

**Conclusion:** Cardiac arrhythmias associate with dialytic phase. Regular dialytic schedule preserves physiological circadian rhythm, but the second day without dialysis is characterized by parasympathetic withdrawal and a steady increase in sympathetic predominance.

**Subject Terms:** Arrhythmias, Autonomic Nervous System, Electrocardiology (ECG), Treatment.

## Introduction

Sudden cardiac death (SCD) is the most common cause of death in end-stage kidney disease (ESKD) patients receiving renal replacement therapy.^1, 2^ Both tachy-and bradyarrhythmias may play a causative role in SCD in ESKD patients on dialysis.^3-5^ Importantly, the dialytic cycle relates to the rate of SCD.^6-9^ Mortality is increased after the long interdialytic interval, and within 6 hours after the end of a hemodialysis session.^6-8^ The dialysis procedure itself triggers multiple mechanisms that can increase the propensity to cardiac arrhythmias.^10^ Oscillations in electrolyte levels, and fluid volume, a pro-inflammatory state, repetitive myocardial injury from uremia and other toxins^10^, and myocardial ‘stunning’^*11*^ are dialysis-specific risk factors of SCD.

A SCD event exemplifies a “perfect storm,” requiring both a susceptible substrate and a trigger.^12^ Potential interactions between fluctuating dialytic cycles – related risk factors of SCD, and traditional SCD triggers can together explain the extremely high rate of SCD in dialysis patients. Autonomic imbalance of the heart is traditionally associated with SCD. Under normal conditions, the autonomic system controls heart rate and rhythm via a balance between the parasympathetic and sympathetic systems. Heart rate variability (HRV) is a measure of fluctuations in the autonomic system and baroreflex sensitivity.^13, 14^ Parasympathetic withdrawal and increased sympathetic input decreases HRV and triggers SCD.^12^ Previous studies using 24-hour Holter ECG recordings reported the association of depressed HRV^15^ with increased mortality in ESKD patients.^16-18^ However, 24-or even 48-hour Holter ECG recordings are too short to study the association of fluctuating autonomic tone with the standard 7-day dialytic cycle. Whether autonomic imbalance associates with both the dialytic cycle and cardiac arrhythmias in dialysis patients remains unknown. Longitudinal changes in autonomic tone before, during, and after dialysis procedures, along with their association with brady-and tachyarrhythmias have not been previously studied. To address this knowledge gap, we conducted an ancillary prospective study in a subset of incident hemodialysis patients, enrolled in the Predictors of Arrhythmic and Cardiovascular Risk in End Stage Renal Disease (PACE) study.^2, 19^ We hypothesized that (1) dialytic cycle is associated with brady-and tachyarrhythmias, and that (2) circadian and dialytic cycles, clinical characteristics, and paroxysmal arrhythmias are associated with periodic changes in heart rate and HRV in incident dialysis patients.

## Methods

### Study population

We conducted a prospective ancillary study within PACE.^19^ Both the parental PACE study and this ancillary study were approved by the Johns Hopkins Institutional Review Board, and all participants provided written informed consent.

The PACE study design has been previously described.^2, 19^ Briefly, the study enrolled adult ESKD patients starting hemodialysis within the six months before enrollment. Patients on home hemodialysis, peritoneal dialysis, in hospice or skilled nursing facility, and patients with an implanted pacemaker or cardioverter-defibrillator were not eligible. At enrollment, participants underwent comprehensive cardiovascular evaluation, which included several types of electrocardiogram (ECG), echocardiogram, cardiac computed tomography, and angiography.

This ancillary study included randomly selected PACE participants who underwent baseline cardiovascular evaluation and agreed to undergo continuous ECG monitoring for at least seven days.

### Clinical covariates

Prevalent coronary artery disease (CAD), cerebrovascular disease (CVD), congestive heart failure (CHF), hypercholesterolemia, hypertension, and diabetes were determined by participants self-report and physician’s diagnosis recorded in the medical record. Left ventricular hypertrophy (LVH) was defined as an echocardiographic left ventricular mass index ≥116 g/m^2^ in males or ≥ 104 g/m^2^ in females. To assess subjective post-dialysis recovery time, participants answered the question: “How long does it take you to recover from a dialysis session?” during a telephone interview, conducted within 30 days of ECG monitoring.

### Continuous ECG monitoring using ECG patch

Continuous ECG monitoring was performed using an ECG patch (Zio Patch, iRhythm Technologies, Inc., San Francisco, CA, USA). During the study visit, a study coordinator applied the device over the left pectorial region,^20^ and instructed the participant to activate a trigger button in the event of cardiac symptoms (presumed arrhythmia). Participants were instructed to wear the adhesive ECG patch for as long as possible, with the goal to obtain at least seven days of continuous ECG recording. After completion of the ECG recording, participants mailed the ECG patch to iRhythm Technologies, Inc, which provided their standard FDA-approved report to the study investigators, via a secure website. The Zio Patch report was reviewed within 24 hours by the study investigators, and clinically important findings were communicated with participants and their healthcare providers. In addition, continuously recorded raw digital ECG signal was provided by iRhythm Technologies, Inc for further analysis.

#### Diagnosis of clinically important arrhythmias

The following clinically important arrhythmias were diagnosed: atrial fibrillation (AF) or flutter (>4 beats), supraventricular tachycardia (SVT, >4 beats), pause >3 seconds, atrioventricular (AV) block of the second or the third degree, ventricular tachycardia (VT, >4 beats), or polymorphic VT / ventricular fibrillation. All arrhythmic events captured by the Zio Patch report were reviewed and validated by at least two study investigators (LGT, RSP, SH).

### Dialytic cycle phases

To determine the association of the dialytic cycle with heart rate and HRV time-series, we categorized the phases of the dialytic cycle relative to their proximity to a dialysis treatment (Figure 1). Every study participant experienced dialysis (4-5 hours), post-dialysis (6 hours immediately after dialysis), between-dialysis (variable length), and pre-dialysis (6 hours preceding dialysis) phases. Over the course of the 7-day cycle, dialysis treatment-adherent study participants were dialyzed every other day for five days (either Monday/Wednesday/Friday or Tuesday/Thursday/Saturday; a total of 3 treatments) and then experienced a two-day long interdialytic interval. For example, for an adherent Monday-Friday schedule, the regular interdialytic period was 32 hours, and the length of the prolonged interdialytic period over the weekend was 56 hours. Non-adherent participants did not follow this standard dialytic routine. Their treatment schedule was interrupted by missed dialysis treatments, leading to a variable ultra-long interdialytic period of at least 72 hours or longer.

**Figure 1.**
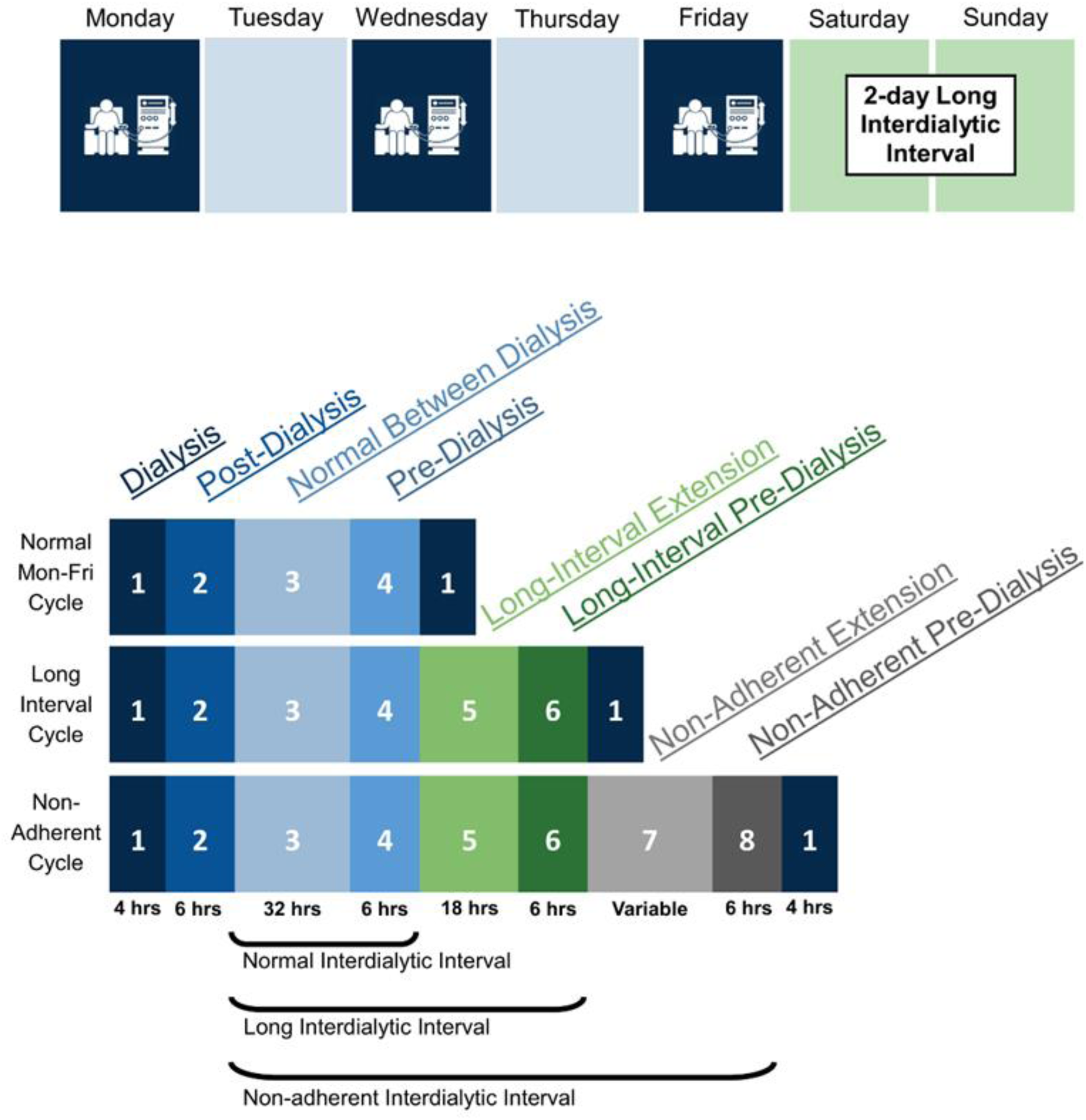
A schematic presentation of the dialytic cycles used for analysis. On top is a representative weekly dialysis schedule for the adherent subjects in this study. Note that some patients may have undergone treatments on Tuesday, Thursday, and Saturday instead of Monday, Wednesday, and Friday. The bottom figure illustrates how phases were broken up for each condition—adherent patients receiving treatment every second day (Mon-Fri), adherent patients during the long two-day interval, and non-adherent patients who missed one or more dialysis sessions. Blue color indicates phases within 48 hours of the last treatment, green color highlights the phases included to analyze the extra 24 hours between treatments in the long interdialytic interval, and grey color symbolizes the phases that were included to analyze >72 hour interdialytic intervals for those that were non-adherent.

### Heart rate variability measurements

A raw single-lead digital ECG signal (sampling rate 200Hz; amplitude resolution 4.88µV) was analyzed in the Tereshchenko laboratory at the Oregon Health & Science University. A custom MATLAB (The MathWorks, Inc, Natick, MA, USA) software application was developed (NMR, EAPA, YLP, MMK; provided at https://github.com/Tereshchenkolab/HRV) to automatically detect QRS complexes and select a single 3-minute normal sinus rhythm epoch for each hour of recording. The algorithm automatically eliminated epochs with premature R_2_beats if the R_1_R_2_interval was shorter than the preceding R_0_R_1_interval by 15% or greater. The algorithm similarly eliminated epochs with a sudden pause, if subsequent the R_1_R_2_interval was longer than the preceding R_0_R_1_interval by 15% or greater, to remove epochs with blocked premature atrial or His extrasystoles, or intermittent sinoatrial or AV block. Traditionally, in Holter ECG analysis, the premature atrial beat was defined by a coupling interval of less than 80% of the mean RR interval.^21^ We applied a more stringent threshold after manually reviewing our ECG data with the thresholds ranging from 2-20%. A sliding 3-minute window approach was used to scan the entirety of the data: when a premature beat (or sudden pause) was detected, the premature beat and subsequent compensatory pause were skipped, and a new 3-minute window search started thereafter again. If the algorithm did not find a continuous 3-minute epoch of sinus rhythm in an entire hour, the software would change the R-peak detection algorithm^22^ and repeated all described above steps. The first R-peak detection algorithm employed was a Pan-Tompkins,^23^ followed by principle component analysis,^24^ and then parabolic fitting.^25^ Because the magnitude of R and S peaks varied within and between patients, the dominant peak of the QRS complex varied during long-term ECG monitoring. We paid special attention to ensure consistent signs of the dominant QRS peak for the entire 3-minute epoch. The greatest average Manhattan distance from baseline to the highest positive (R) peak and highest negative (S) peak was calculated to identify the best dominant peak for each 3-minute epoch. The accuracy of consistent dominant (R or S) peaks detection, and accuracy of the selection of consecutive normal sinus beat were validated on a data subset by the investigator (NMR), with the aid of a graphical display (Figure 2).

**Figure 2.**
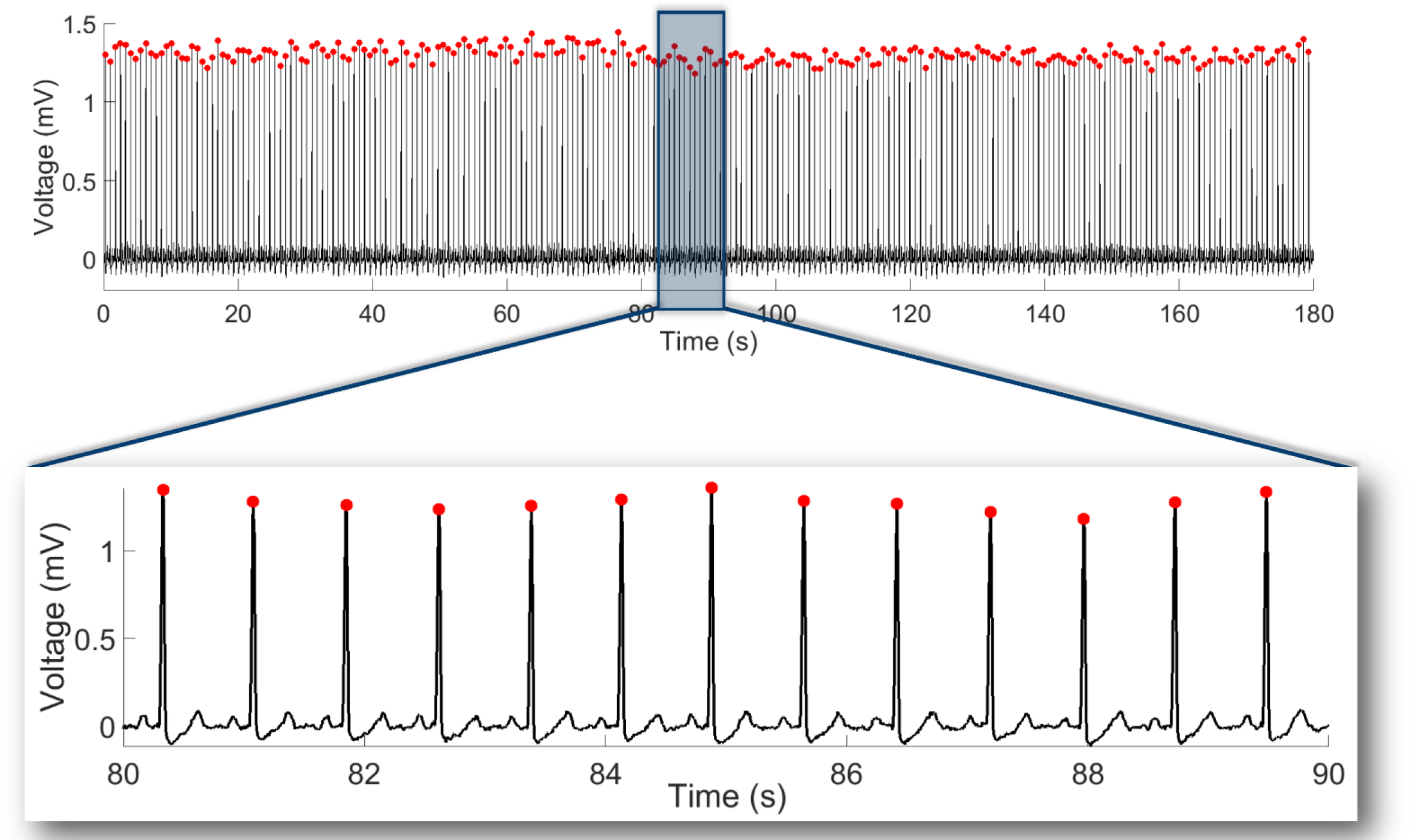
A representative example of a single-lead ECG with detected R-peaks and measured RR’ intervals. A three-minute epoch is shown, with mean heart rate 78 bpm, rMSSD of 11.5 ms, LF power of 2.4 s^2^, HF power of 6.3 s^2^, LF/HF ratio of 0.38, SD_1_of 8.2 ms, SD_2_of 19.7 ms, SD_12_ratio of 0.41, sample entropy of 2.0, and Renyi entropy of 1.4. A ten-second portion is displayed for closer examination of ECG morphology.

HRV was measured according to the Standards.^26, 27^ Developed (by MMK) MATLAB (the MathWorks, Inc, Natick, MA, USA) software code is provided at https://github.com/Tereshchenkolab/HRV.

#### Time-domain heart rate variability measures

Heart rate and the root mean square of the successive normal sinus to normal sinus (NN) intervals differences (rMSSD) were calculated for each 3-minute epoch selected per hour.

#### Frequency-domain heart rate variability measures

The low frequency (LF; 0.04-0.15 Hz) power, high frequency (HF; 0.15-0.4 Hz) power, and LF/HF ratio of powers were calculated for each 3-minute epoch.

#### Nonlinear heart rate variability measures

Quantitative analysis of the Poincaré plot was performed.^27^ The Poincaré plot was derived from every 3-minute NN data epoch by plotting the values NN_n+1_ against the values of NN_n_. SD_1_was calculated as the standard deviation (SD) of the cloud of points in the direction perpendicular to the line-of-identity. SD_2_was calculated as the SD of the cloud of points in the direction of the line-of-identity. SD_1_/SD_2_ratio was called SD_12_.

#### Entropy

To quantify the entropy rate on a short-length NN series, we elected to measure sample entropy^27^ and Renyi entropy for each 3-minute epoch. Renyi entropy was calculated as described by Cornforth *et al*.^28^ We used an α value equal to 4 because of previous data suggesting that positive α (1-5) provides the best discrimination of cardiac autonomic neuropathy, and based on the sampling rate of our data.^29^

### Statistical analysis

Normality of the distribution of continuous variables was evaluated using standardized normal probability (P-P) plot. For comparison of clinical characteristics in participants with versus without detected arrhythmia, normally distributed continuous variables were presented as means ± SD and were compared using a *t*-test. Fisher’s exact test was used to compare categorical variables.

The main dataset was structured as a panel of time-series, as HRV was measured at the beginning of each hour (assuming equal intervals between 3-minute epochs). The assumption of equal intervals between 3-min epochs was confirmed for 81% of epochs starting in the first second of each hour (44%), within first 10 minutes (76%) or first 15 minutes (81%) of each hour. The assumption of equal intervals between 3-min epochs was violated for 10% of epochs starting in the second half of an hour. To test the robustness of our findings, we conducted sensitivity analysis after exclusion of epochs that violated equal intervals assumption required for time-series analysis.

Within-subject and between-subjects SDs were reported for each phase of dialysis. Paired comparison of HRV in different phases of the dialytic cycle was performed using analysis of variance within subjects (for repeated measures). As previous studies reported an increased risk of SCD in post-dialysis phase after the long interdialytic weekend,^30^ we performed paired comparisons of HRV in the post-dialysis phase that followed (1) normal dialytic cycle vs. (2) two-day long interdialytic interval vs. (3) longer than 72-hour interval without dialysis in non-adherent participants.

To determine whether demographic and clinical characteristics, paroxysmal cardiac arrhythmias, and dialytic and circadian (24-hour) cycles are associated with time-series of heart rate and HRV metrics, we constructed autoregressive conditional heteroscedasticity (ARCH)/ generalized autoregressive conditionally heteroscedastic (GARCH) panel models.^31^ In ARCH time series analysis, both mean and variance of HRV metric were modeled as time-dependent, which allowed modeling volatility that can arise in response to the dialysis procedure or other unmeasured factors. Time series of heart rate and HRV metrics (one-by-one) served as an outcome. To identify our ARCH/GARCH model, we first explored autocorrelation function (ACF) and partial autocorrelation (PACF) function of the HRV time-series, and ACF/PACF of squared HRV variables values. As ACF and PACF of heart rate and HRV time-series represented white noise, but ACF/PACF of squared series tapered (autoregressive of order 1), we constructed ARCH(1/1)/GARCH(1) models.

To determine whether demographic and clinical characteristics are associated with heart rate and HRV time-series, age, sex, race, prevalent CAD, CHF, CVD, history of AF, diabetes, Charlson comorbidity index (CCI), and post-dialysis recovery time were included in each ARCH model.

To determine whether paroxysmal cardiac arrhythmias are associated with time-series of heart rate and HRV metrics, separate ARCH models were constructed for each type of incident arrhythmia (VT, AF, SVT, pause). ARCH models were adjusted for age, sex, race, prevalent CAD, CHF, CVD, AF, CCI, and post-dialysis recovery time.

To describe the circadian (24-hour) rhythm, while accounting for multiple 24-day cycles analyzed for the same patient (longitudinal/panel data structure), we constructed periodic regression models with fixed (within) estimator. We used periodic regression to analyze a behavior of heart rate and HRV that vary in a circular-scale 24-hour cycle. We converted the circular variable “hour in a 24-hour cycle” from cyclic (hours of the day) to angular (radians) format, and then to trigonometric format (paired units sine and cosine). We studied whether heart rate and HRV time-series responded in a periodic way to 24-hour cycle. We estimated the periodic mean (mesor) for heart rate and HRV metrics, and calculated an amplitude (A) of variation about the mesor in the modeled cycle:

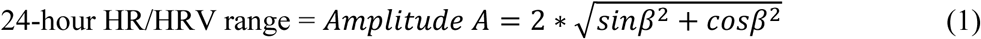

We calculated phase angle (acrophase φ) as a time within the 24-hour cycle when heart rate/HRV is maximized (peak location): φ = arctan(sinβ/cosβ). The minimum is located 12 hours (0.5 cycles) away from the maximum. As dialysis and the immediate post-dialytic phase are well-known to significantly affect heart rate and HRV,^30, 32^ dialysis (phase 1) and post-dialytic phase 2 were excluded from periodic regression analyses. We stratified circadian rhythm analyses by the type of interdialytic phase: in a regular dialytic schedule (phases 3-4), second-day interdialytic extension (phases 5-6), and interdialytic extension above 72 hours (phases 7-8 in non-adherent participants missing dialysis), as shown in Figure 1.

To determine whether the dialytic cycle is associated with the heart rate and HRV time-series after removal of the effect of circadian rhythm, we constructed periodic panel ARCH/GARCH models to determine a change in HRV per hour of remoteness from the 1st dialysis hour. ARCH/GARCH models were adjusted for age, sex, race, prevalent CAD, CHF, CVD, AF, diabetes, CCI, and post-dialysis recovery time. In addition, circadian 24-hour cycle (paired units sine and cosine) were included in each ARCH model and served to adjust for circadian periodicity.

#### Sensitivity analysis

To test the robustness of our findings, we excluded 10% of epochs that violated the equal intervals assumption that is required for time-series ARCH models.

Statistical analysis was performed using STATA MP 15.1 (StataCorp LP, College Station, TX).

## Results

### Study population

The study population (Table 1) included 28 PACE participants (mean age 54±13 y; 57% men; 96% black). Approximately one-third of the population had a history of structural heart disease with normal left ventricular ejection fraction (LVEF; 70±9%). The average dialysis recovery time was 16 minutes.

**Table 1.** Clinical characteristics of study participants.

| Characteristic | Total (n=28) | With<br>arrhythmia<br>(n=13) | Arrhythmia-free (n=15) | P-value |
| --- | --- | --- | --- | --- |
| Age(SD), y | 53.9(12.5) | 60.8(8.3) | 47.9(12.7) | 0.004 |
| Male, n(%) | 16(57.1) | 8(61.5) | 8(53.3) | 0.718 |
| Black, n(%) | 27(96.4) | 12(92.3) | 15(100.0) | 0.464 |
| Body mass index(SD), kg/m<sup>2</sup> | 29.4(7.4) | 29.0(8.7) | 29.7(6.4) | 0.835 |
| Body surface area(SD), m<sup>2</sup> | 2.00(0.23) | 1.98(0.25) | 2.02(0.21) | 0.668 |
| Cause of ESKD: |  |  |  |  |
| Glomerulonephritis, n(%) | 3(10.7) | 2(15.4) | 1(6.7) | 1.00 |
| Hypertension, n(%) | 5(17.9) | 2(15.4) | 3(20.0) | 1.00 |
| Diabetes, n(%) | 12(42.9) | 6(46.2) | 6(40.0) | 1.00 |
| HIV/Other/Genetic/Obstruction, n(%) | 7(25.0) | 3(23.1) | 4(26.7) | 1.00 |
| Unknown, n(%) | 1(3.6) | 0 | 1(6.7) | 1.00 |
| Hypertension, n(%) | 28(100) | 13(100.0) | 15(100) | 1.00 |
| Diabetes, n(%) | 16(57.1) | 9(69.2) | 7(46.7) | 0.276 |
| Coronary Artery Disease, n(%) | 8(28.6) | 6(46.2) | 2(13.3) | 0.096 |
| Heart failure, n(%) | 9(32.1) | 5(38.5) | 4(26.7) | 0.689 |
| Cerebrovascular disease, n(%) | 8(28.6) | 5(38.5) | 3(20.0) | 0.410 |
| History of atrial fibrillation, n(%) | 10(35.7) | 5(38.5) | 5(33.3) | 1.00 |
| Hypercholesterolemia, n(%) | 19(67.9) | 10(76.9) | 9(60.0) | 0.435 |
| Smoker current or former, n(%) | 15(53.6) | 7(53.8) | 8(53.3) | 0.885 |
| Drinker current or former, n(%) | 23(82.1) | 12(92.3) | 11(73.3) | 0.325 |
| Charlson comorbidity index(SD) | 5.5(2.2) | 5.8(1.9) | 5.1(2.2) | 0.391 |
| LV Ejection Fraction(SD), % | 70.3(9.3) | 68.8(9.6) | 71.7(9.0) | 0.441 |
| LV Internal Dimension Diastole(SD), cm | 5.3(0.7) | 5.5(0.9) | 5.2(0.4) | 0.309 |
| LV Internal Dimension Systole(SD), cm | 3.1(0.7) | 3.3(0.9) | 3.0(0.5) | 0.317 |
| LV mass index(SD), g/m<sup>2</sup> | 146.0(53.2) | 164.2(66.7) | 130.6(34.0) | 0.152 |
| LV Hypertrophy, n(%) | 17(70.8) | 9(81.8) | 8(61.5) | 0.386 |
| 3-mo Averaged calcium (SD), mg/dl | 8.52(0.65) | 8.54(0.58) | 8.50(0.73) | 0.868 |
| 3-mo Averaged potassium (SD), mEq/L | 4.34(0.39) | 4.45(0.41) | 4.24(0.35) | 0.177 |
| 3-mo Averaged intradialytic weight<br>change (SD), kg | 2.09(0.97) | 2.14(1.20) | 2.06(0.83) | 0.869 |
| Dialysis recovery time(SD), min | 15.9(3.3) | 19.6(6.6) | 12.7(2.1) | 0.335 |

ESKD=end-stage kidney disease. 3-month-averaged values are from the start of dialysis.

### Cardiac arrhythmia events

Almost half of the participants (n=13, 46%) had arrhythmias detected during monitoring (Table 2). Except for one patient with 46% paroxysmal AF burden, all detected events were non-sustained (NS), lasting < 30 second, and asymptomatic. NS VT was more frequent during dialysis or within 6 hours post-dialysis, as compared to pre-or between-dialysis (63% vs. 37%, P=0.015). SVT occurred more frequently pre-or between-dialysis as compared to during-or post-dialysis (84% vs. 16%, P=0.015). All patients with NS VT were free from CAD at baseline. No events of high degree AV block were detected. Participants with detected arrhythmias were older than arrhythmia-free individuals, but no other significant differences were observed (Table 1).

**Table 2.** Cardiac arrhythmia events timeline.

|  | Events 6h-pre-dialysis, n (%) | Events during dialysis, n(%) | Events 6h-post-dialysis, n(%) | Events between-dialysis, n(%) | Number of patients (% of population) |
| --- | --- | --- | --- | --- | --- |
| Total number of events, n=47 | 12(26) | 8(17) | 9(19) | 18(38) | 13(46) |
| VT, n=8 | 1(13) | 2(25) | 3(38) | 2(25) | 4(15) |
| SVT, n=17 | 5(29) | 1(6) | 2(12) | 9(53) | 9(32) |
| AF, n=21 | 6(29) | 4(19) | 4(19) | 7(33) | 2(7) |
| Pauses>3 s, n=1 | 0 | 1(100) | 0 | 0 | 1(3.5) |

VT=ventricular tachycardia; SVT=supraventricular tachycardia; AF=atrial fibrillation

### Dynamic time series of heart rate and heart rate variability

The average duration of analyzed continuous ECG recording was 6.5±2.3 days. All 28 participants had at least one ECG recording over a two-day long interdialytic interval, and four participants missed 1-2 dialysis sessions (up to 120 hours) during ECG monitoring.

Average heart rate was the fastest within the first 6 hours post-dialysis (Table 3) and afterward gradually slowed down (Figure 3). Differences in heart rate among participants were greater than differences in heart rate in the same participant during ECG monitoring.

**Table 3.** Heart rate and heart rate variability in different phases of dialytic cycle.

| HRV measure | Phase 1<br>N=323;<br>n=26<br>T=12.4 | Phase 2<br>N=480;<br>n=26<br>T=18.5 | Phase 3<br>N=2554;<br>n=28<br>T=91.2 | Phase 4<br>N=438;<br>n=27<br>T=16.2 | Phase 5<br>N=428;<br>n=28<br>T=15.3 | Phase 6<br>N=135;<br>n=26<br>T=5.2 | Phase 7<br>N=221;<br>n=4<br>T=55.3 | Phase 8<br>N=15;<br>n=3<br>T=5 | Within<br>ANOVA<br>P |
| --- | --- | --- | --- | --- | --- | --- | --- | --- | --- |
| Heart rate(SD), bpm | 74.1(11.3) | 81.3(13.4) | 77.2(12.2) | 75.9(11.5) | 77.6(14.1) | 73.9(12.2) | 70.8(8.0) | 71.0(5.4) | <0.0001 |
| SD between | 10.4 | 11.3 | 10.1 | 10.3 | 12.9 | 13.2 | 9.8 | 5.3 |  |
| SD within | 6.1 | 7.6 | 7.7 | 6.0 | 6.0 | 4.4 | 4.3 | 3.1 |  |
| rMSSD(SD), ms | 12.6(7.8) | 9.7(5.2) | 11.0(6.4) | 11.1(6.7) | 11.1(5.6) | 11.8(5.6) | 11.3(4.1) | 9.2(2.3) | <0.0001 |
| SD between | 6.1 | 3.7 | 4.1 | 5.2 | 3.7 | 4.3 | 2.7 | 2.1 |  |
| SD within | 4.6 | 3.7 | 5.1 | 5.1 | 4.4 | 4.2 | 3.6 | 1.7 |  |
| HF power(SD), s<sup>2</sup> | 4.6(2.0) | 4.1(1.8) | 4.3(1.9) | 4.4(2.0) | 4.5(2.0) | 4.6(2.2) | 5.8(2.2) | 5.1(2.9) | <0.0001 |
| SD between | 1.5 | 1.3 | 1.4 | 1.5 | 1.5 | 1.7 | 1.9 | 2.9 |  |
| SD within | 1.3 | 1.4 | 1.4 | 1.4 | 1.3 | 1.4 | 1.4 | 0.9 |  |
| LF power(SD), s<sup>2</sup> | 3.1(1.0) | 3.0(1.0) | 3.1(1.0) | 2.9(0.9) | 3.0(1.0) | 3.0(1.0) | 3.1(1.0) | 3.0(0.8) | 0.068 |
| SD between | 0.6 | 0.6 | 0.6 | 0.5 | 0.6 | 0.7 | 0.8 | 0.3 |  |
| SD within | 0.8 | 0.8 | 0.8 | 0.8 | 0.8 | 0.8 | 0.8 | 0.7 |  |
| LF/HF ratio(SD) | 0.82(0.46) | 0.89(0.48) | 0.87(0.46) | 0.84(0.45) | 0.84(0.47) | 0.80(0.43) | 0.66(0.39) | 0.88(0.58) | 0.006 |
| SD between | 0.36 | 0.36 | 0.35 | 0.37 | 0.36 | 0.33 | 0.37 | 0.61 |  |
| SD within | 0.27 | 0.31 | 0.32 | 0.32 | 0.29 | 0.26 | 0.24 | 0.16 |  |
| Poincaré SD<sub>1</sub>(SD), ms | 9.0(5.6) | 6.9(3.7) | 7.8(4.6) | 7.9(4.8) | 7.8(4.0) | 8.4(3.9) | 8.0(2.9) | 6.5(1.6) | <0.0001 |
| SD between | 4.3 | 2.6 | 2.9 | 3.7 | 2.6 | 3.0 | 1.9 | 1.5 |  |
| SD within | 3.3 | 2.6 | 3.6 | 3.6 | 3.1 | 2.9 | 2.6 | 1.2 |  |
| Poincaré SD<sub>2</sub>(SD), ms | 25.8(18.7) | 21.6(15.9) | 23.8(18.4) | 23.7(17.2) | 22.4(16.5) | 24.4(16.5) | 20.3(15.3) | 23.5(17.0) | 0.002 |
| SD between | 15.2 | 11.1 | 14.9 | 19.6 | 12.1 | 12.5 | 13.7 | 16.6 |  |
| SD within | 11.1 | 11.6 | 13.5 | 11.4 | 12.2 | 10.7 | 10.1 | 7.4 |  |
| SD<sub>12</sub> (SD), % | 0.44(0.25) | 0.42(0.25) | 0.43(0.24) | 0.43(0.27) | 0.46(0.28) | 0.45(0.27) | 0.56(0.33) | 0.47(0.32) | 0.461 |
| SD between | 0.20 | 0.19 | 0.19 | 0.20 | 0.20 | 0.20 | 0.34 | 0.33 |  |
| SD within | 0.16 | 0.17 | 0.18 | 0.20 | 0.19 | 0.18 | 0.20 | 0.12 |  |
| Sample Entropy(SD) | 1.49(0.40) | 1.39(0.46) | 1.39(0.44) | 1.43(0.44) | 1.42(0.44) | 1.43(0.47) | 1.52(0.43) | 1.25(0.62) | 0.0004 |
| SD between | 0.25 | 0.26 | 0.21 | 0.24 | 0.23 | 0.29 | 0.23 | 0.73 |  |
| SD within | 0.35 | 0.40 | 0.39 | 0.38 | 0.37 | 0.38 | 0.38 | 0.30 |  |
| Renyi entropy(SD) | 1.31(0.31) | 1.33(0.28) | 1.31(0.30) | 1.33(0.29) | 1.31(0.31) | 1.29(0.31) | 1.23(0.33) | 1.30(0.36) | 0.758 |
| SD between | 0.17 | 0.13 | 0.09 | 0.17 | 0.12 | 0.15 | 0.13 | 0.31 |  |
| SD within | 0.27 | 0.26 | 0.29 | 0.27 | 0.29 | 0.28 | 0.31 | 0.24 |  |

N=number of observations; n=number of participants; T=average number of hours; SD=standard deviation; SD1=Poincaré plot
standard deviation perpendicular the line of identity; SD2=Poincaré plot standard deviation along the line of identity.

**Figure 3.**
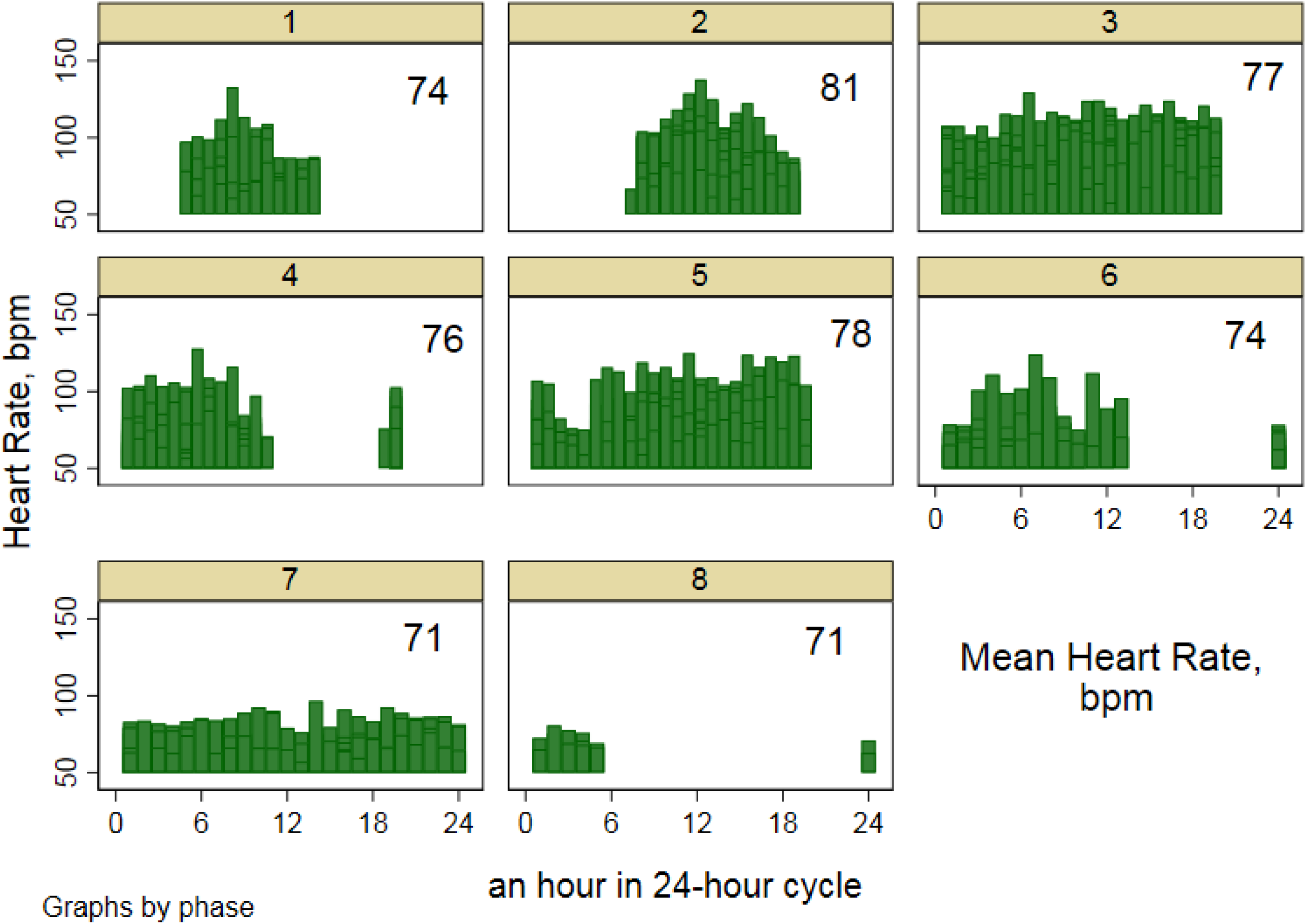
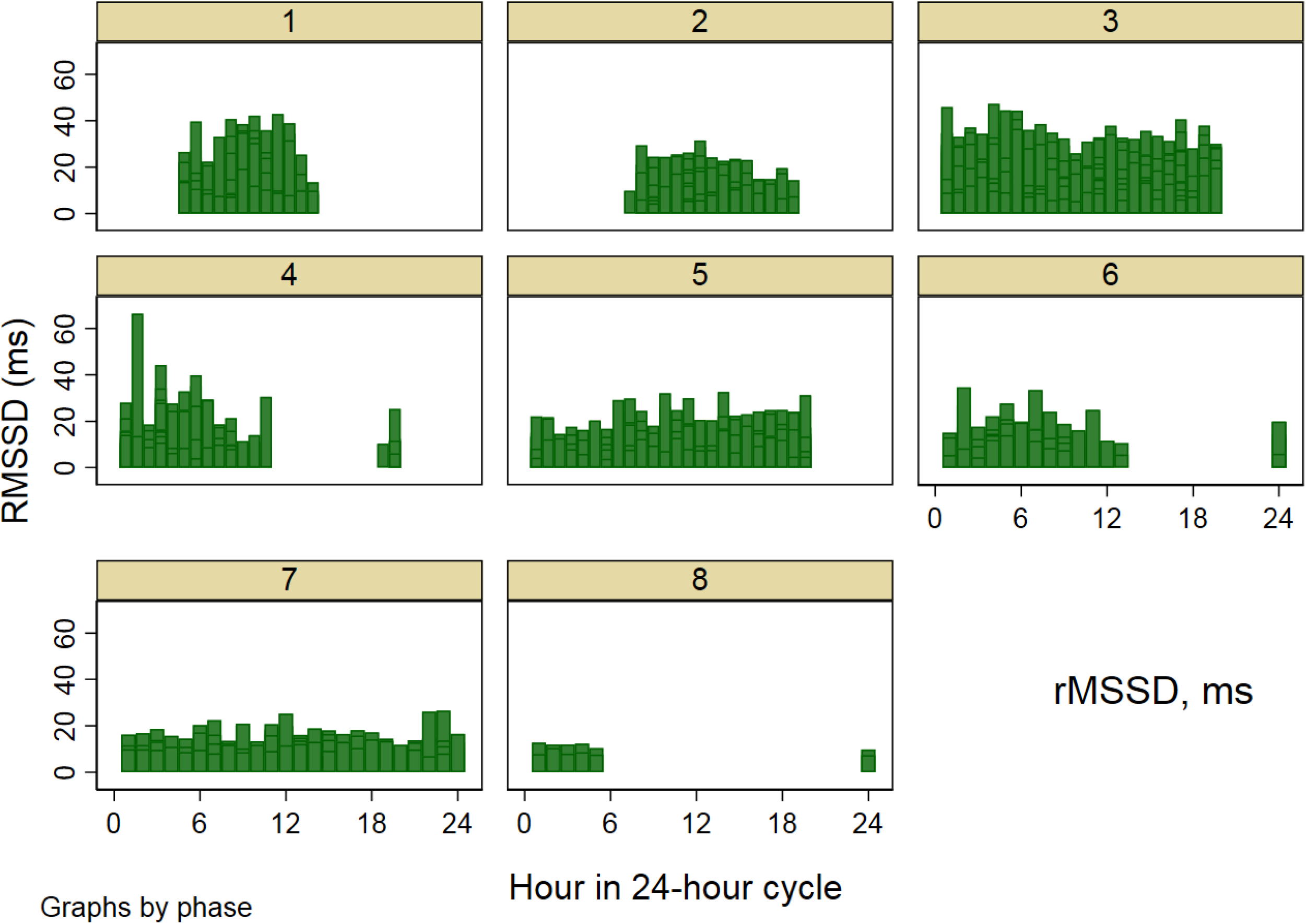
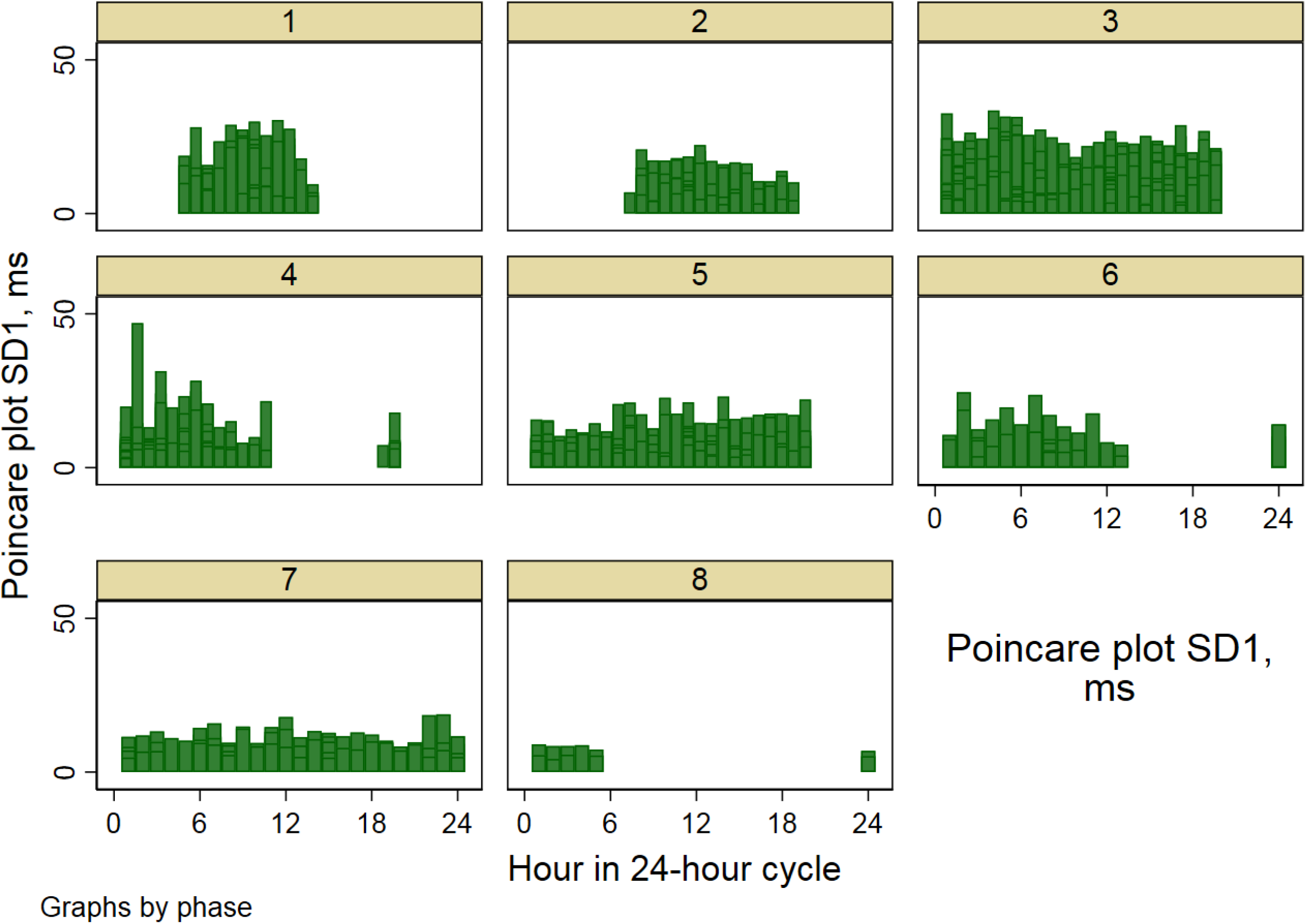
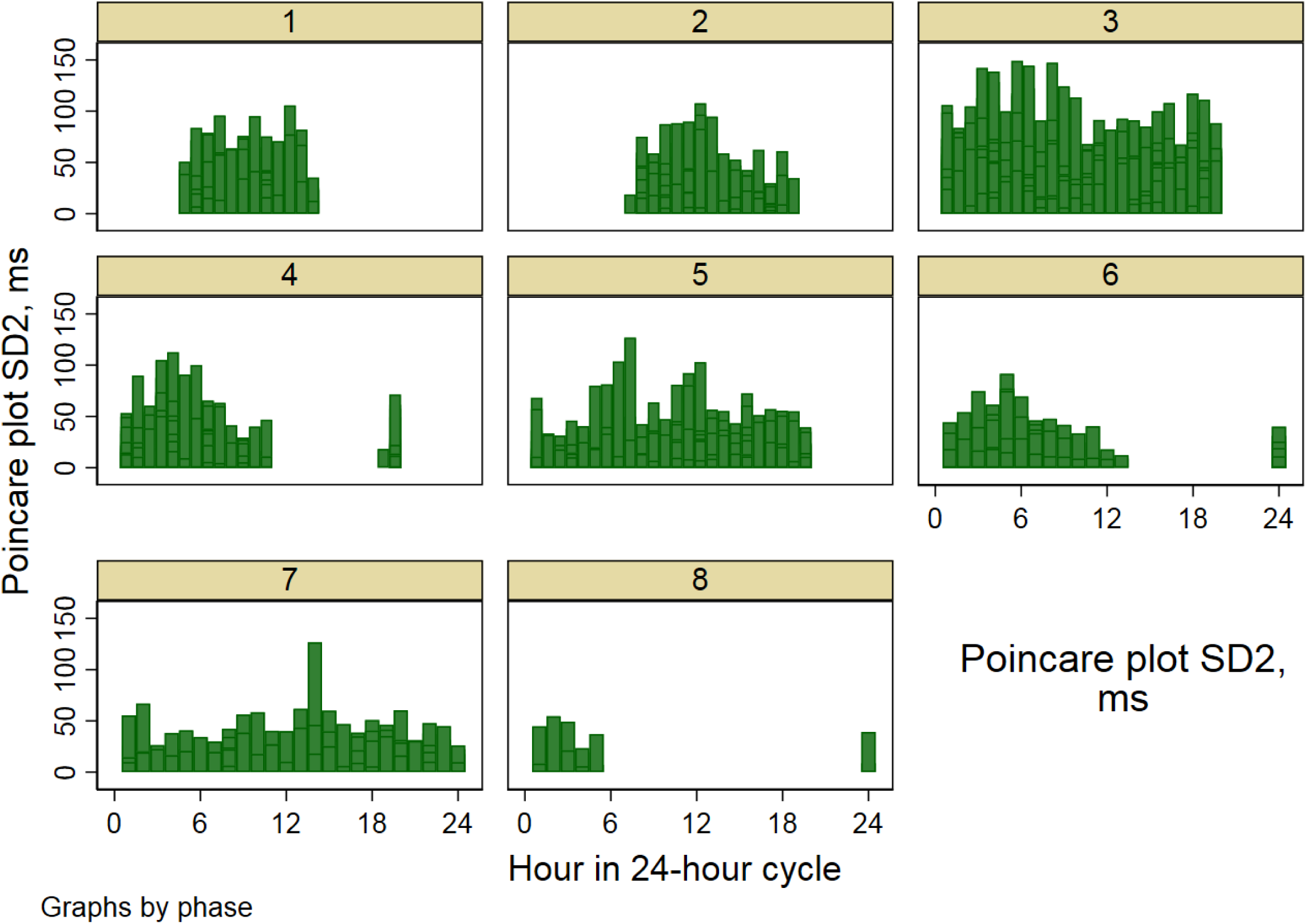
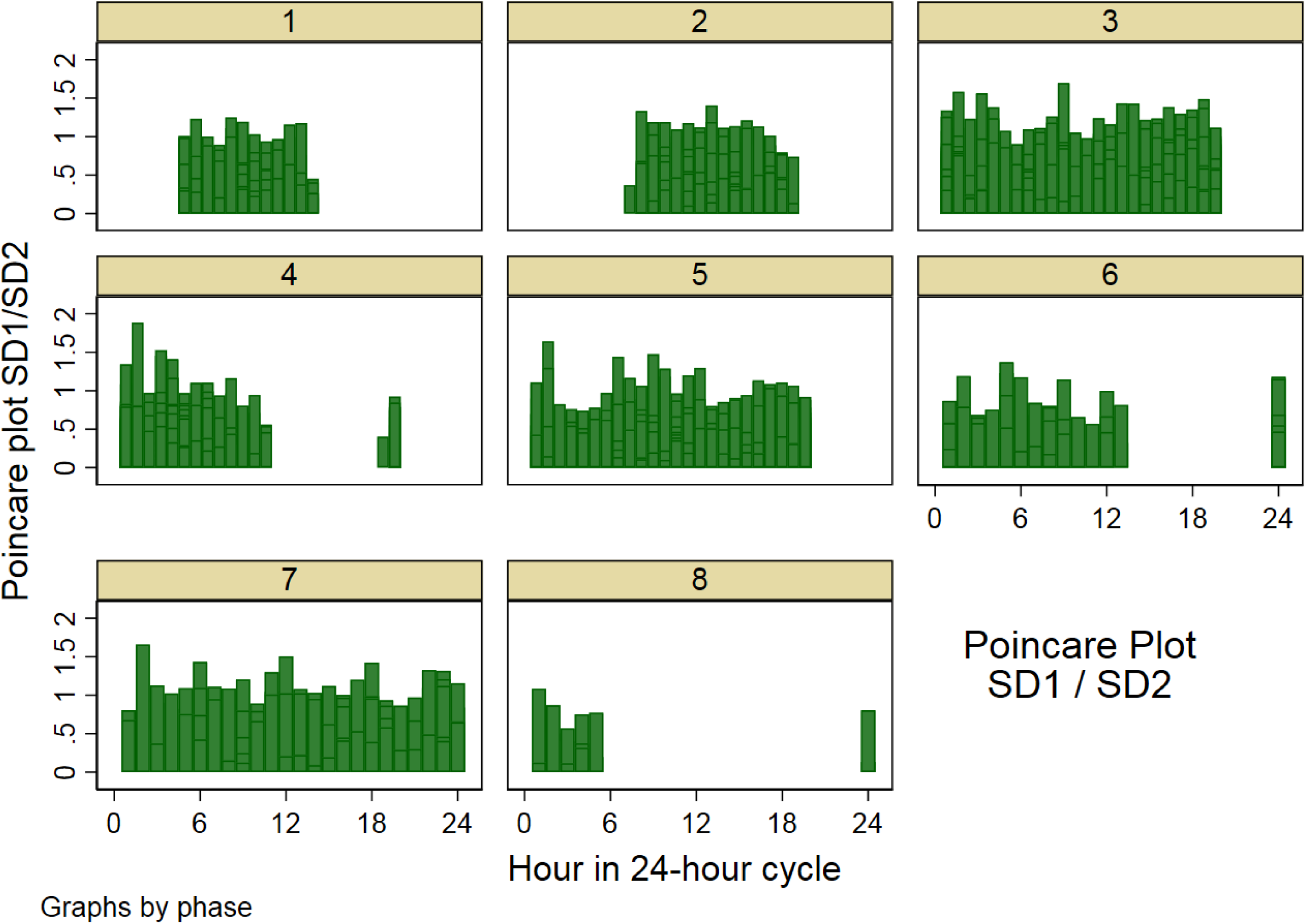
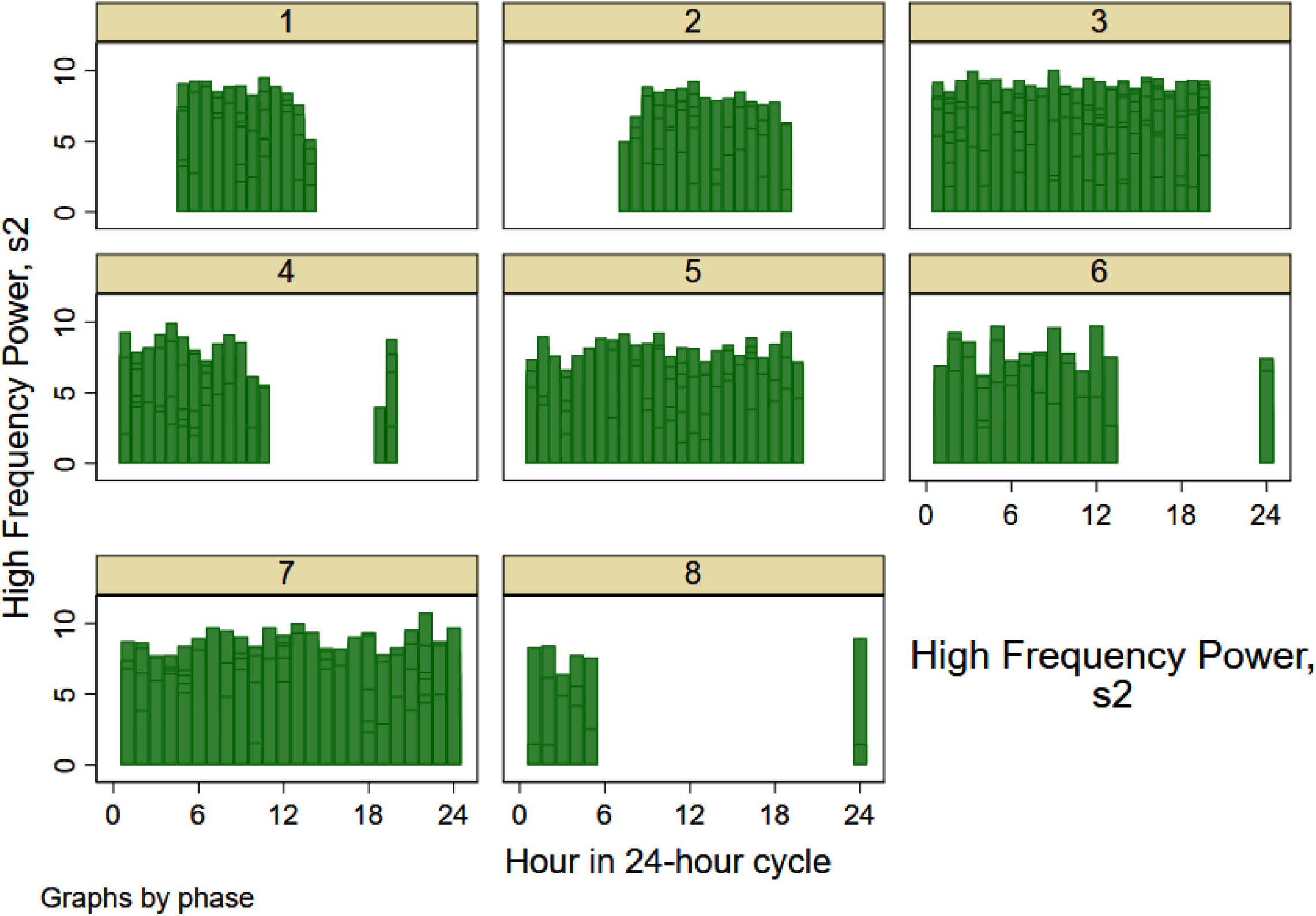
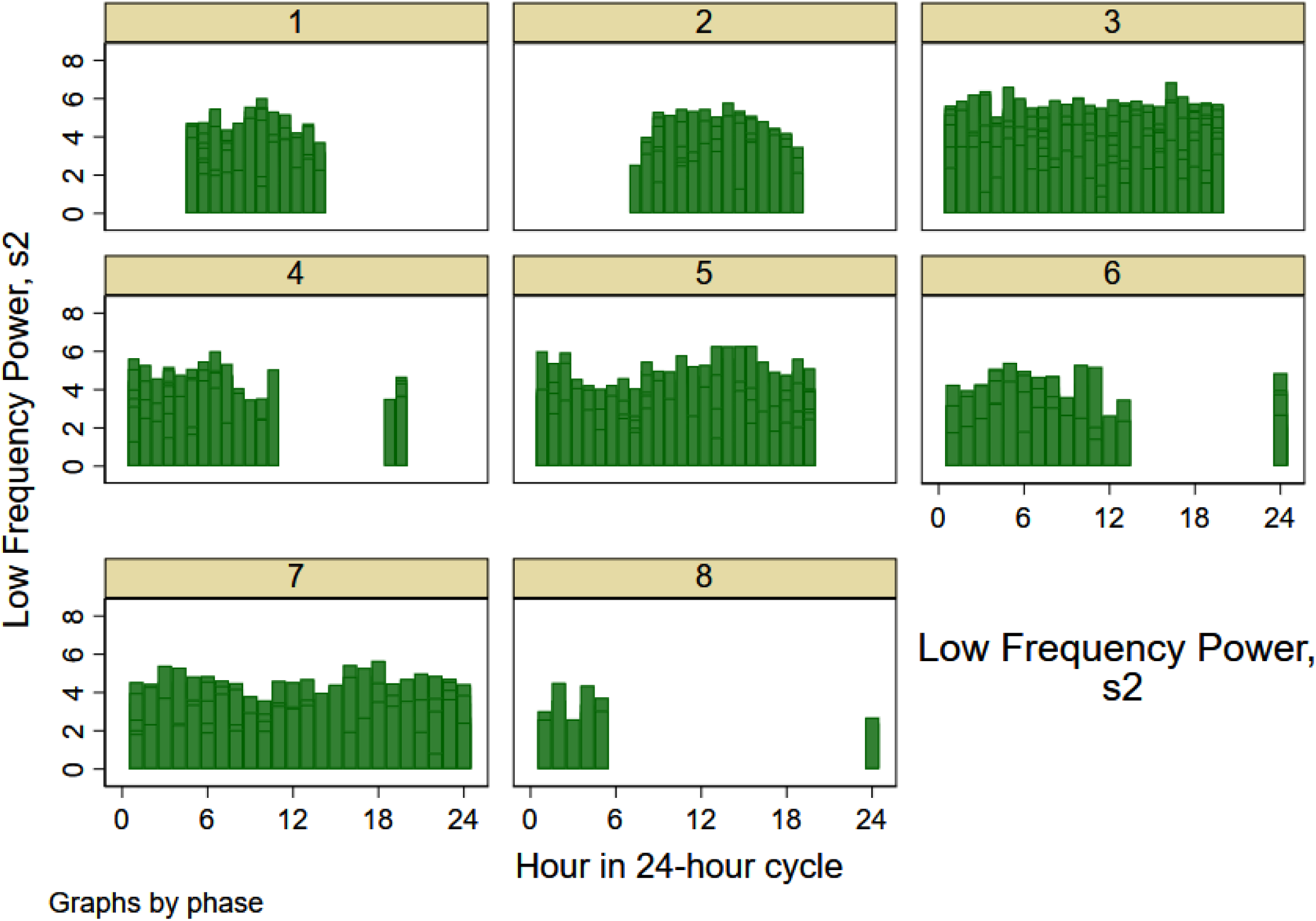
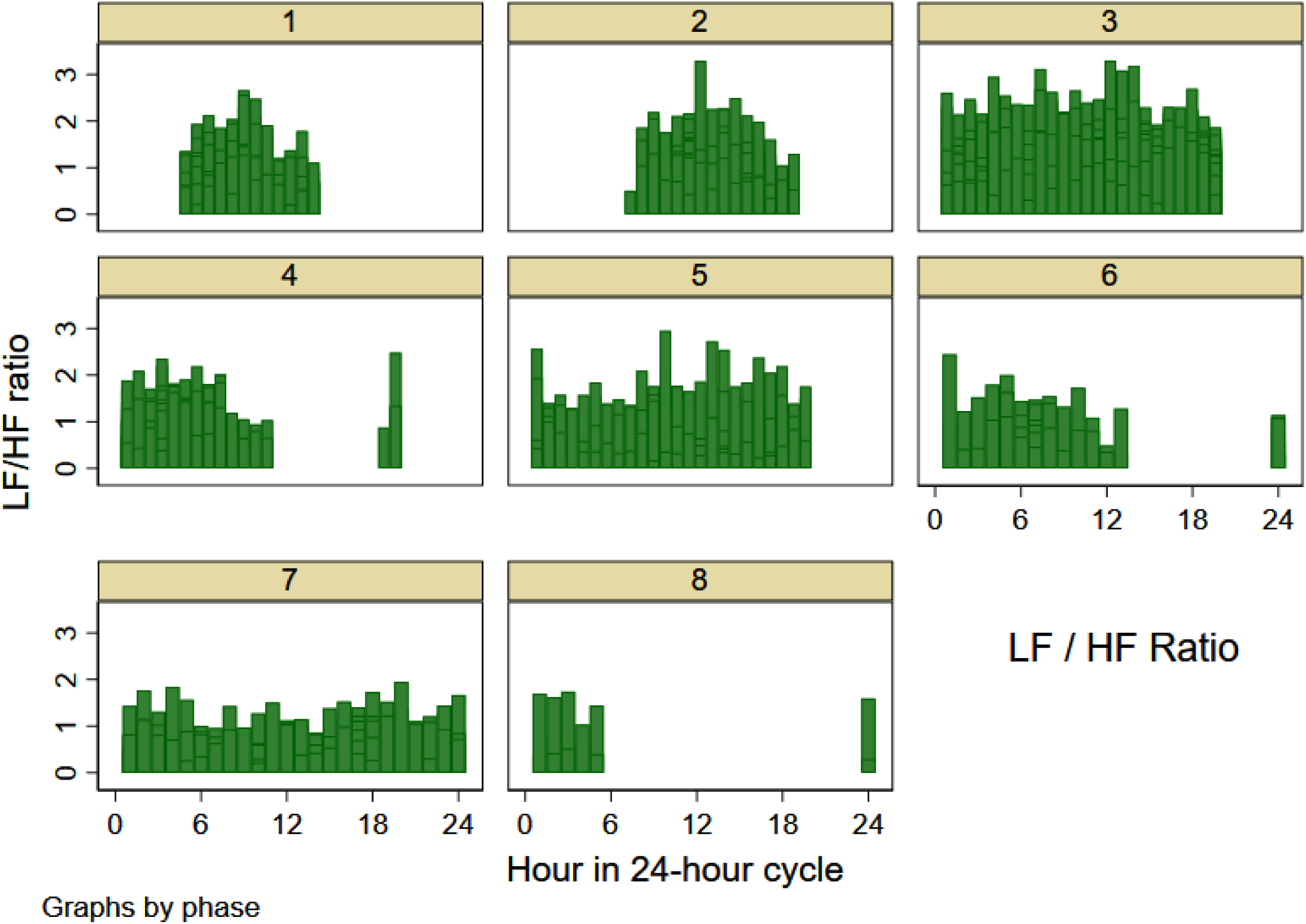
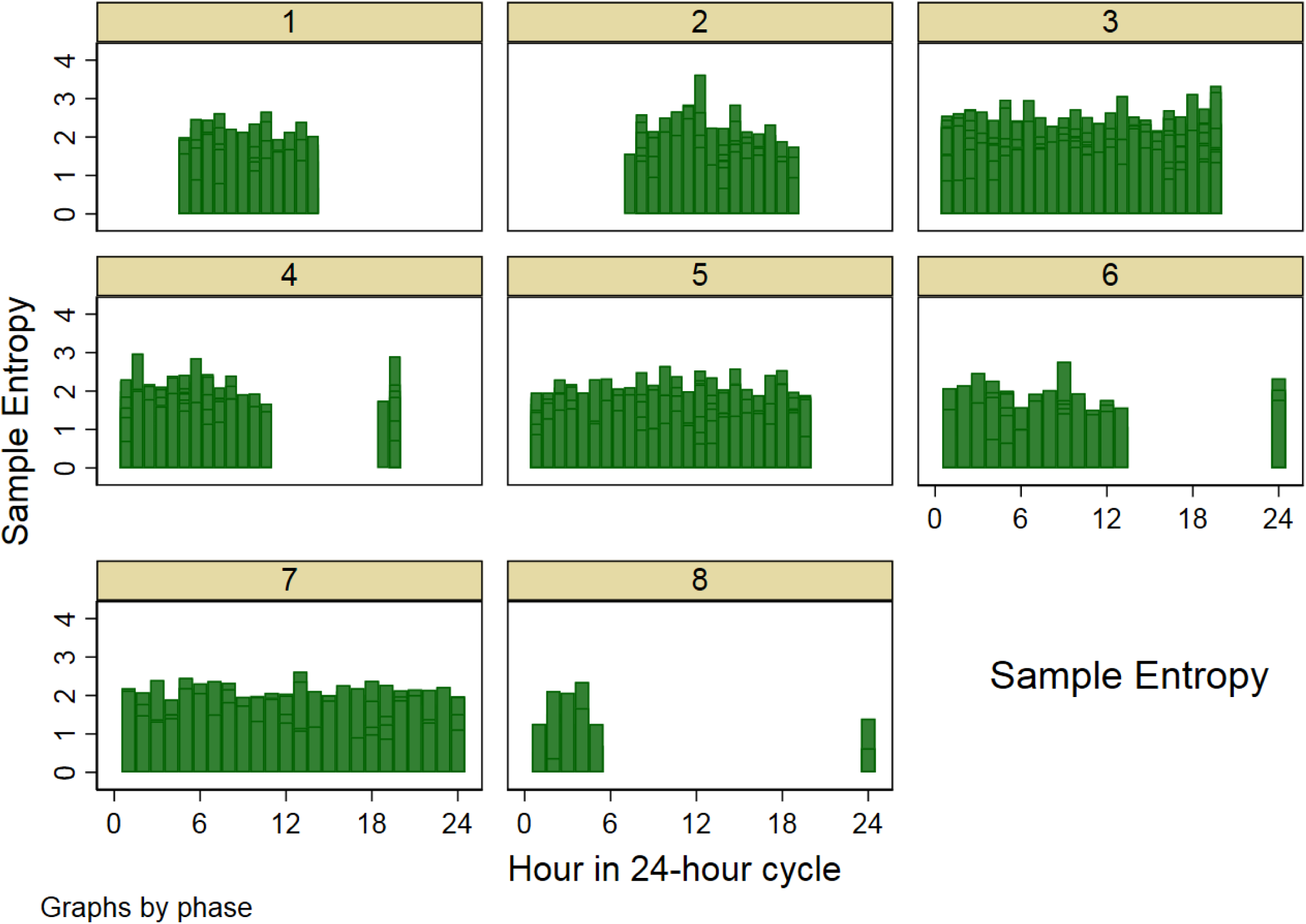
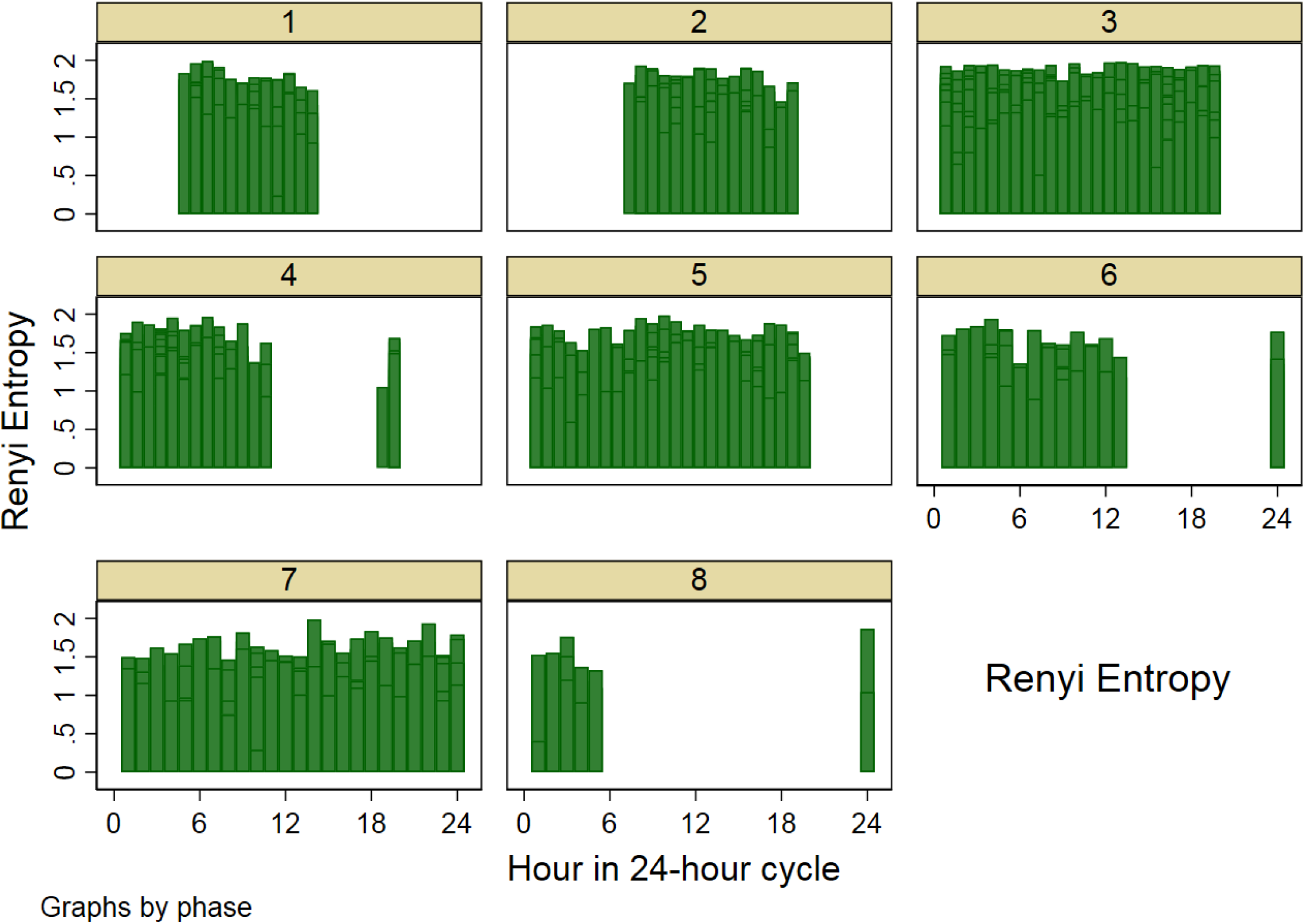
Panel-data bar plot of (**A**) heart rate, (**B**) rMSSD, (**C**) Poincaré SD_1_, (**D**) Poincaré SD_2_, (**E**) Poincaré SD_12_, (**F**) High frequency (HF) power, (**G**) Low frequency (LF) power, (**H**) LF/HF ratio, (I) Sample entropy, (J) Renyi Entropy plotted versus 24-hour cycle for each phase of the dialytic cycle.

Short-term HRV (rMSSD, HF power, Poincaré SD_1_) decreased post-dialysis, then recovered by the time of the next regular dialysis session. However, if dialysis was not performed within the next 72 hours, short-term HRV gradually diminished further (Table 3). Differences in HRV among participants were comparable with differences in HRV in the same participant over time.

In contrast, the greatest differences in entropy were observed within each individual participant, highlighting the importance of patient-specific entropy changes over time. Differences in entropy between participants were small, reflecting a similar stage of the disease.

In an unadjusted paired comparison, there were no statistically significant differences in HRV post-dialysis after different durations of the preceding interdialytic interval (Table 4).

**Table 4.**
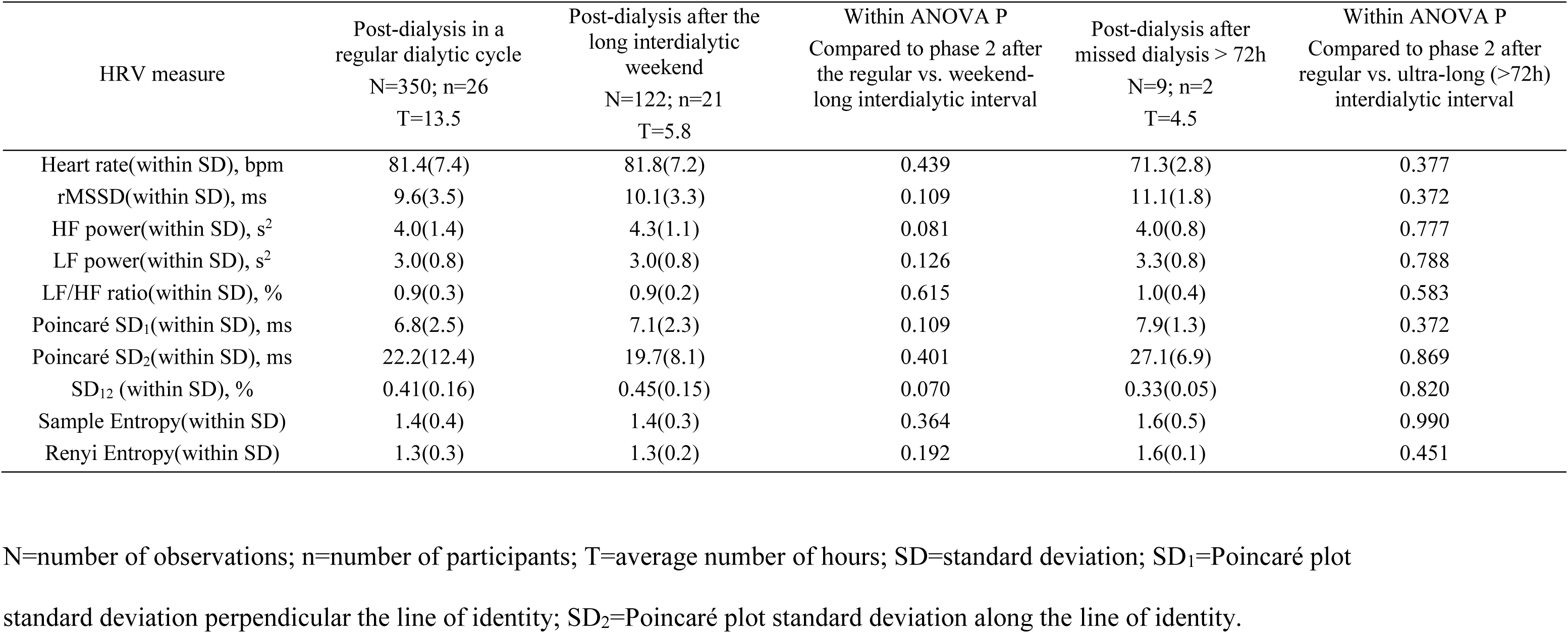
Comparison of heart rate and HRV post-dialysis in a regular dialytic (every other day) cycle, after two-day long interdialytic interval, and after missed dialysis for more than 72 hours.

### Association of demographic and cardiovascular risk factors with heart rate and HRV in different types of dialytic cycles

During the regular (every other day) dialytic cycle, prevalent cardiovascular disease, and its risk factors were associated with faster heart rate and more depressed HRV (Table 5), with the exception of AF history. History of AF was associated with slower heart rate and relatively preserved HRV, perhaps due to the use of beta-blockers or calcium channel blockers. There was no association of CAD, CHF, and CCI with entropy.

**Table 5A.** Association of demographic and clinical characteristics with heart rate and short-term HRV in ARCH models, stratified by the type of dialytic cycle.

| Clinical characteristic |  | Heart rate (95%CI), bpm | P | rMSSD (95%CI), ms | P | SD1 (95%CI), ms | P | HF power(95%CI), s<sup>2</sup> | P |
| --- | --- | --- | --- | --- | --- | --- | --- | --- | --- |
| Regular dialytic cycle<br>(phases 1-4) | Age, /10y | -2.9(-3.1 to-2.7) | <0.0001 | +0.9(0.8-1.0) | <0.0001 | +0.7(0.6-0.7) | <0.0001 | +0.4(0.3-0.5) | <0.0001 |
|  | Female sex | +8.1(7.5-8.6) | <0.0001 | +0.7(0.4-0.9) | <0.0001 | +0.5(0.3-0.6) | <0.0001 | +0.05(-0.07-0.17) | 0.437 |
|  | Black race | +23(-388-433) | 0.927 | -2(-21 to 17) | 0.825 | -2(-15 to 12) | 0.824 | -7(-21 to 6) | 0.301 |
|  | CAD | +4.5(3.8-5.3) | <0.0001 | +0.03(-0.3 to 0.4) | 0.849 | +0.02(-0.2 to 0.3) | 0.846 | -2.1(-2.2 to -1.9) | <0.0001 |
|  | CVD | +6.8(6.1-7.5) | <0.0001 | -2.7(-3.1 to -2.4) | <0.0001 | -1.9(-2.2 to -1.7) | <0.0001 | -0.5(-0.7 to -0.3) | <0.0001 |
|  | CHF | +2.3(1.5-3.1) | <0.0001 | -2.4(-2.7 to -2.0) | <0.0001 | -1.7(-1.9 to -1.4) | <0.0001 | +1.4(1.2-1.6) | <0.0001 |
|  | AF history | -10.5(-11.0 to-10.1) | <0.0001 | +5.3(5.1-5.6) | <0.0001 | +3.8(3.6-4.0) | <0.0001 | +0.6(0.5-0.8) | <0.0001 |
|  | Diabetes | +1.1(0.6-1.5) | <0.0001 | -1.8(-2.0 to -1.5) | <0.0001 | -1.3(-1.4 to -1.1) | <0.0001 | +1.3(1.2-1.4) | <0.0001 |
|  | CCI | +1.2(1.1-1.3) | <0.0001 | -0.5(-0.6 to -0.4) | <0.0001 | -0.4(-0.03 to-0.01) | <0.0001 | -0.10(-0.13 to -0.07) | <0.0001 |
|  | Recovery time, min | +0.04(0.01-0.6) | 0.012 | +0.03(-0.05 to -0.01) | <0.0001 | -0.02(-0.03 to -0.01) | <0.0001 | -0.05(-0.06 to -0.04) | <0.0001 |
| 2<sup>nd</sup> day interdialytic extension<br>(phases 5-6) | Age, /10y | -4.0(-4.4 to-3.7) | <0.0001 | -0.2(-0.4-0.1) | 0.265 | -1.0(-0.3 to 0.1) | 0.274 | +0.3(0.2-0.5) | <0.0001 |
|  | Female sex | +3.8(2.8-4.8) | <0.0001 | -1.5(-2.1 to -0.8) | <0.0001 | -1.0(-1.5 to -0.6) | <0.0001 | +0.07(-0.24 to0.38) | 0.656 |
|  | Black race | -15(-21 to -9)) | <0.0001 | -8(-13 to -3) | 0.003 | -5(-9 to -2) | 0.004 | -7.5(-9.8 to -5.1) | <0.0001 |
|  | CAD | -5.3(-6.7 to -4.0) | <0.0001 | +0.06(-0.9 to 1.0) | 0.906 | +0.04(-0.6 to 0.7) | 0.903 | -2.2(-2.8 to -1.7) | <0.0001 |
|  | CVD | +6.1(4.4-7.7) | <0.0001 | -1.3(-2.3 to -0.4) | 0.007 | -0.9(-1.6 to -0.3) | 0.006 | -0.7(-1.2 to -0.2) | 0.004 |
|  | CHF | +4.5(2.8-6.2) | <0.0001 | -2.2(-3.3 to -1.2) | <0.0001 | -1.6(-2.3 to -0.9) | <0.0001 | +1.2(0.6-1.8) | <0.0001 |
|  | AF history | -2.3(-3.4 to -1.2) | <0.0001 | +4.0(3.3-4.7) | <0.0001 | +2.8(2.4-3.3) | <0.0001 | +1.2(0.9-1.6) | <0.0001 |
|  | Diabetes | -2.4(-0.4 to 0.6) | <0.0001 | +0.5(-0.2 to 1.1) | 0.173 | +0.3(-0.1 to 0.8) | 0.176 | +1.9(1.6-2.2) | <0.0001 |
|  | CCI | +0.25(0.05-0.60) | 0.100 | -0.3(-0.5 to -0.2) | <0.0001 | -0.2(-0.3 to -0.1) | <0.0001 | -0.05(-0.12 to 0.03) | 0.199 |
|  | Recovery time, min | -0.23(-0.29 to-0.17) | <0.0001 | -0.05(-0.10 to-0.0007) | 0.047 | -0.03(-0.07 to -0.0002) | 0.048 | -0.06(-0.08 to -0.04) | <0.0001 |
| Phases 7-8 | Age, /10y | -5.3(-5.8 to-4.7) | <0.0001 | -0.20(-0.24 to -0.15) | <0.0001 | -1.4(-1.7 to -1.0) | <0.0001 | +0.5(0.3-0.7) | <0.0001 |
|  | Diabetes | -4.8(-6.0 to-3.6) | <0.0001 | -1.6(-2.5 to-0.6) | 0.001 | -1.10(-1.8 to -0.4) | 0.001 | +2.2(1.8-2.6) | <0.0001 |
|  | Recovery time, min | +2.4(2.2-2.6) | <0.0001 | +0.4(0.13-0.64) | 0.003 | +0.3(0.09-0.5) | 0.003 | +0.3(0.2-0.5) | <0.0001 |

**Table 5B.** Association of demographic and clinical characteristics with intermediate HRV in ARCH models, stratified by the type of dialytic cycle.

| Clinical characteristic |  | LF power(95%CI),<br>s<sup>2</sup> | P | LF/HF ratio<br>(95%CI) | P | SD1/SD2 ratio<br>(95%CI) | P | SD2 (95%CI), ms | P |
| --- | --- | --- | --- | --- | --- | --- | --- | --- | --- |
| Regular dialytic cycle<br>(phases 1-4) | Age, /10y | -0.1(-1.0 to -0.04) | <0.0001 | -0.08(-0.09 to -0.07) | <0.0001 | +0.03(0.03-0.04) | <0.0001 | -0.14(-0.17 to -0.11) | <0.0001 |
|  | Female sex | +0.17(0.10-0.24) | <0.0001 | -0.05(-0.07 to -0.02) | <0.0001 | +0.02(0.01-0.04) | <0.0001 | -0.3(-0.9 to 0.2) | 0.190 |
|  | Black race | +3.3(1.0-5.7) | 0.005 | +2.5(-1.2 to 6.2) | 0.182 | -0.9(-2.3-0.4) | 0.177 | +32(-121 to 187) | 0.678 |
|  | CAD | +0.6(0.5-0.7) | <0.0001 | +0.64(0.61-0.68) | <0.0001 | -0.2(-0.2 to -0.21) | <0.0001 | +7.0(5.9-8.1) | <0.0001 |
|  | CVD | -0.15(-0.26 to -0.05) | 0.005 | -0.05(-0.08 to -0.01) | 0.009 | -0.03(-0.05 to -0.01) | 0.002 | -1.8(-2.6 to -1.0) | <0.0001 |
|  | CHF | -0.62(-0.74 to -0.50) | <0.0001 | -0.52(-0.56 to -0.48) | <0.0001 | +0.19(0.17-0.21) | <0.0001 | -3.1(-4.3 to -1.9) | <0.0001 |
|  | AF history | -0.07(-0.15 to 0.01) | 0.102 | -0.23(-0.26 to -0.21) | <0.0001 | +0.03(0.02-0.05) | <0.0001 | +5.3(4.7-5.9) | <0.0001 |
|  | Diabetes | -0.59(-0.67 to -0.52) | <0.0001 | -0.40(-0.42 to -0.38) | <0.0001 | 0.20(0.19-0.22) | <0.0001 | -19.4(-20.1 to -18.7) | <0.0001 |
|  | CCI | +0.02(0.01-0.04) | 0.011 | +0.05(0.04-0.05) | <0.0001 | -0.01(-0.01to-0.01) | <0.0001 | -0.9(-1.0 to -0.7) | <0.0001 |
|  | Recovery time, min | +0.03(0.02-0.03) | <0.0001 | +0.021(0.019-0.22) | <0.0001 | -0.01(-0.01to-0.01) | <0.0001 | +0.32(0.29-0.35) | <0.0001 |
| 2<sup>nd</sup> day interdialytic extension<br>(phases 5-6) | Age, /10y | +0.04(-0.03-0.1) | 0.265 | -0.06(-0.08 to -0.03) | <0.0001 | +0.01(-0.004-0.03) | 0.139 | -1.0(-2.0 to -1.0) | <0.0001 |
|  | Female sex | +0.13(-0.05 to 0.31) | 0.170 | -0.03(-0.10 to 0.03) | 0.296 | +0.03(-0.008-0.06) | 0.133 | -1.3(-2.7 to 0.5) | 0.058 |
|  | Black race | +3.0(1.5-4.5) | <0.0001 | +1.8(1.4-2.2) | <0.0001 | -1.0(-1.3 to -0.7) | <0.0001 | +37(26-47) | <0.0001 |
|  | CAD | +0.5(0.2-0.7) | <0.0001 | +0.51(0.43-0.59) | <0.0001 | -0.2(-0.3 to -0.2) | <0.0001 | +8.2(6.0-10.5) | <0.0001 |
|  | CVD | -0.4(-0.7 to -0.2) | 0.001 | +0.001(-0.09-0.10) | 0.983 | -0.03(-0.1-0.04) | 0.426 | -3.4(-6.1 to -0.7) | 0.015 |
|  | CHF | -0.7(-1.0 to -0.4) | <0.0001 | -0.32(-0.42 to -0.22) | <0.0001 | +0.19(0.12-0.25) | <0.0001 | -4.5(-7.2 to -1.8) | 0.001 |
|  | AF history | -0.2(-0.4 to -0.02) | 0.027 | -0.2(-0.3 to -0.1) | <0.0001 | +0.07(0.03-0.12) | 0.001 | +4.5(2.3-6.7) | <0.0001 |
|  | Diabetes | -0.7(-0.9 to -0.6) | <0.0001 | -0.5(-0.6 to -0.4) | <0.0001 | +0.22(0.18-0.26) | <0.0001 | -20(-22 to -19) | <0.0001 |
|  | CCI | +0.03(0.02-0.05) | 0.276 | +0.02(0.01-0.04) | <0.0001 | -0.004(-0.1-0.01) | 0.930 | -0.6(-1.1 to -0.2) | 0.009 |
|  | Recovery time, min | +0.03(0.02-0.05) | <0.0001 | +0.02(0.01-0.02) | <0.0001 | -0.008(-0.01 to -0.005) | <0.0001 | +0.4(0.3-0.5) | <0.0001 |
| Phases 7-8 | Age, /10y | +0.2(0.1-0.3) | 0.001 | -0.01(-0.10-0.03) | 0.740 | -0.02(-0.05 to -0.001) | 0.041 | -0.8(-2.0 to 0.1) | 0.100 |
|  | Diabetes | -1.4(-1.6 to -1.2) | <0.0001 | -0.47(-0.53 to -0.41) | <0.0001 | +0.43(0.39-0.47) | <0.0001 | -14.6(-16.0 to -13.3) | <0.0001 |
|  | Recovery time, min | +0.11(0.06-0.16) | <0.0001 | -0.08(-0.09 to -0.06) | <0.0001 | +0.03(0.02-0.04) | <0.0001 | -4.3(-4.7 to -3.8) | <0.0001 |

**Table 5C.** Association of demographic and clinical characteristics with entropy in ARCH models, stratified by the type of dialytic cycle.

|  | Clinical characteristic | Sample entropy (95%CI), bpm | P | Renyi entropy (95%CI), ms | P |
| --- | --- | --- | --- | --- | --- |
| Regular dialytic cycle<br>(phases 1-4) | Age, per 10y | +0.06(0.05-0.08) | <0.0001 | -0.007(-0.02 to 0.003) | 0.189 |
|  | Female sex | +0.08(0.05-0.11) | <0.0001 | -0.03(-0.06 to -0.001) | 0.006 |
|  | Black race | +0.3(-0.6 to 1.2) | 0.530 | -0.19(-0.43 to 0.06) | 0.138 |
|  | CAD | +0.0004(-0.05 to 0.05) | 0.988 | +0.0005(-0.03 to 0.03) | 0.974 |
|  | CVD | -0.18(-0.23 to -0.14) | <0.0001 | +0.0001(-0.03 to 0.03) | 0.992 |
|  | CHF | +0.01(-0.04 to 0.06) | 0.709 | +0.02(-0.02 to 0.06) | 0.244 |
|  | AF history | +0.11(0.07-0.14) | <0.0001 | +0.09(0.06-0.12) | <0.0001 |
|  | Diabetes | -0.10(-0.13 to -0.07) | <0.0001 | -0.07(-0.09 to -0.04) | <0.0001 |
|  | CCI | -0.002(-0.01 to 0.005) | 0.567 | -0.006(-0.01 to 0.0001) | 0.052 |
|  | Recovery time, min | +0.002(-0.0001 to 0.004) | 0.051 | -0.003(-0.005 to -0.001) | <0.0001 |
| 2<sup>nd</sup> day interdialytic extension<br>(phases 5-6) | Age, per 10y | +0.05(0.02-0.09) | <0.0001 | -0.003(-0.003 to 0.02) | 0.832 |
|  | Female sex | -0.01(-0.06 to 0.09) | 0.713 | -0.06(-0.11 to -0.001) | 0.045 |
|  | Black race | +0.4(-0.2 to 1.0) | 0.200 | +0.03(-0.4 to 0.4) | 0.914 |
|  | CAD | +0.13(0.02-0.24) | 0.017 | +0.09(0.005-0.18) | 0.037 |
|  | CVD | -0.24(-0.36 to -0.13) | <0.0001 | -0.10(-0.18 to -0.02) | 0.018 |
|  | CHF | -0.15(-0.28 to -0.02) | 0.022 | -0.05(-0.02 to 0.05) | 0.295 |
|  | AF history | +0.06(-0.03 to 0.14) | 0.193 | +0.11(0.04-0.17) | 0.001 |
|  | Diabetes | +0.02(-0.06 to 0.09) | 0.653 | -0.08(-0.13 to -0.02) | 0.009 |
|  | CCI | +0.01(-0.002 to 0.1) | 0.090 | -0.01(-0.02 to 0.004) | 0.151 |
|  | Recovery time, min | +0.005(-0.0002 to 0.10) | 0.062 | -0.001(-0.005 to 0.003) | 0.733 |
| Phases<br>7-8 | Age, per 1y | -0.003(-0.008 to 0.003) | 0.360 | +0.03(-0.008 to 0.07) | 0.116 |
|  | Diabetes | -0.51(-0.61 to -0.42) | <0.0001 | -0.16(-0.23 to -0.08) | <0.0001 |
|  | Recovery time, min | +0.11(0.09-0.14) | <0.0001 | +0.006(-0.002 to 0.01) | 0.126 |

In contrast, during the second day of the long interdialytic period, traditional cardiovascular risk factors and black race were associated with slower heart rates (Table 5). Female sex, black race, and prevalent CAD were associated with increased SD_2_ and LF power, suggesting greater sympathetic predominance. Interestingly, diabetes was associated with a smaller SD_2_ and LF power, implying that HRV-manifestation of autonomic disbalance can vary.

In our study, four participants (100% black; 2 males and two females) missed dialysis for >72h. They were CAD-and CHF-free, with a short subjective post-dialysis recovery time (9±2 min), but high CCI (6.8±1.2). During missed dialysis phases 7-8, diabetes and longer perceived dialysis recovery time were associated with greater degree of HRV depression (Table 5).

### Association of heart rate and heart rate variability with cardiac arrhythmias

In a fully adjusted ARCH model, significantly increased heart rate [by 11.2 (95%CI 10.1-12.3) bpm (P<0.0001)] was associated with paroxysmal VT events. There was no association of VT events with HRV. There were no associations of other types of observed arrhythmia with heart rate or HRV.

### Circadian rhythm in heart rate and HRV during different types of dialytic cycles

During the interdialytic period in a regular (every other day) dialytic cycle, heart rate, and all HRV parameters demonstrated a significant circadian pattern, as expected (Table 6). Short-term HRV (rMSSD and SD_1_) peaked at night, whereas heart rate and SD_12_ peaked during the day.

**Table 6.** Comparison of circadian rhythm in heart rate and HRV during different interdialytic phases.

| Regular interdialytic period (phases 3-4) |  |  |  |  | 2<sup>nd</sup> day of a long interdialytic interval (phases 5-6) |  |  |  | Nonadherent interdialytic extension (phases 7-8) |  |  |  |
| --- | --- | --- | --- | --- | --- | --- | --- | --- | --- | --- | --- | --- |
| HR / HRV metric | Mesor (95%CI) | 24-hour Amplitude (95%CI) | Peak time (95%CI) | P Sin Cos | Mesor (95%CI) | 24-hour Amplitude (95%CI) | Peak time (95%CI) | P Sin Cos | Mesor (95%CI) | 24-hour Amplitude (95%CI) | Peak time (95%CI) | P Sin Cos |
| HR, bpm | 75.4(74.6-76.3) | 6.8(5.6-8.6) | 07:44(06:52 - 08:14) | <0.0001 | 74.8(72.9-76.7) | 6.9(5.2-11.1) | 07:30(06:08-10:44) | 0.008<br>0.020 | 73.7(71.9-75.6) | 8.3(3.4-13.4) | 15:30(14:15-15:48) | 0.006<br><0.0001 |
| rMSSD, ms | 10.6(0.9-11.2) | 1.5(1.0-3.1) | 02:01(20:22-03:16) | 0.342<br><0.0001 | NS | NS | NS | 0.067<br>0.182 | 8.7(7.2-10.2) | 6.8(2.7-10.9) | 05:05(04:47-06:18) | 0.001<br>0.085 |
| HF, s<sup>2</sup> | 4.6(4.5-4.8) | 1.0(0.6-1.5) | 17:28(17:04-18:23) | <0.0001<br>0.166 | 4.9(4.5-5.3) | 1.1(0.2-2.2) | 16:54(16:36-22:39) | 0.038<br>0.202 | NS | NS | NS | 0.604<br>0.235 |
| LF, s<sup>2</sup> | 3.2(3.1-3.3) | 0.44(0.17-0.72) | 16:50(16:38-17:34) | 0.001<br>0.022 | NS | NS | NS | 0.090<br>0.163 | 3.5(3.1-3.8) | 1.0(0.1-1.9) | 16:16(16:00-16:17) | 0.032<br>0.028 |
| LF/HF | 0.82(0.78-0.85) | 0.14(0.06-0.24) | 05:49(05:11-08:34) | 0.003<br>0.770 | NS | NS | NS | 0.271<br>0.246 | 0.83(0.73-0.93) | 0.42(0.14-0.71) | 16:31(16:26-16:59) | 0.003<br>0.012 |
| SD<sub>1</sub>, ms | 7.5(7.1-7.9) | 1.1(0.7-2.2) | 02:00(20:22-03:15) | 0.343<br>0<0.0001 | NS | NS | NS | 0.067<br>0.182 | 6.2(5.1-7.2) | 4.8(1.9-7.8) | 05:05(04:47-06:18) | 0.001<br>0.085 |
| SD<sub>2</sub>, ms | 21.7(20.2-23.2) | 6.6(2.9-10.8) | 02:58(00:47-03:31) | 0.025<br><0.0001 | 18.7(15.0-22.3) | 11.7(1.9-21.9) | 03:56(01:53-04:06) | 0.032<br>0.007 | NS | NS | NS | 0.067<br>0.784 |
| SD<sub>12</sub> | 0.45(0.43-0.47) | 0.057(0.008-0.12) | 15:54(07:58-16:08) | 0.088<br>0.023 | NS | NS | NS | 0.112<br>0.078 | NS | NS | NS | 0.450<br>0.789 |
| SamEn | 1.37(1.32-1.42) | 0.081(0.077-0.204) | 02:08(18:41-03:30) | 0.478<br>0.014 | NS | NS | NS | 0.877<br>0.825 | NS | NS | NS | 0.631<br>0.117 |
| RenEn | 1.28(1.25-1.31) | 0.11(0.02-0.20) | 05:03(04:43-08:31) | 0.019<br>0.195 | 1.21(1.12-1.30) | 0.25(0.08-0.48) | 05:28(04:54-10:07) | 0.022<br>0.506 | NS | NS | NS | 0.420<br>0.163 |

NS=non-significant (absent circadian rhythm)

The long interdialytic period abolished circadian rhythms, with few exceptions. Periodic changes in heart rate remained largely unchanged across all interdialytic periods, but its acrophase shifted to the afternoon in an ultra-long (missed dialysis > 72 hours) interdialytic interval. On the second day of the long interdialytic period, the amplitude of circadian rhythm in Renyi entropy and SD_2_ increased, whereas circadian rhythm in other HRV metrics dissipated. During an ultra-long intradialytic period in nonadherent participants who missed dialysis for at least 72 hours, we observed distorted periodic pattern in short-term HRV (rMSSD, SD_1_) with acrophase in the afternoon, whereas circadian rhythm in other HRV metrics remained eliminated.

### Association of the dialytic cycle with heart rate and HRV

In fully adjusted ARCH models (Table 7), dialysis and post-dialytic phases 1-2 were characterized by gradual improvement of parasympathetic tone (increasing rMSSD and SD_1_). During the regular interdialytic phase in every other day dialysis cycle (phases 3-4), we observed very few significant trends: slight decrease in rMSSD and SD_1_, and the increase in SD_2_ and sample entropy (Table 7). In contrast, during the second day extension of interdialytic period, there were significant and meaningful trends in all HRV parameters: heart rate gradually increased, short-term HRV (rMSSD, HF power, SD_1_) and SD_12_ decreased, whereas intermediate HRV (SD_2_, LF/HF ratio, Renyi entropy) increased, suggesting depressed parasympathetic and increased sympathetic influences. Missing dialysis for > 72h was associated with a steady increase in SD_12_ ratio and HF power (Table 7).

**Table 7.** Association of the dialytic cycle with heart rate and HRV after adjustment for circadian rhythm, clinical characteristics.

|  | Dialysis & postdialysis (phases 1-2) |  | Regular interdialytic phases 3-4 |  | 2<sup>nd</sup> day interdialytic extension (phases 5-6) |  | Nonadherent interdialytic extension (phases 7-8) |  |
| --- | --- | --- | --- | --- | --- | --- | --- | --- |
| HR / HRV metric | per 24-h remoteness from 1<sup>st</sup> dialysis hour | P | per 24-h remoteness from 1<sup>st</sup> dialysis hour | P | per 24-h remoteness from 1<sup>st</sup> dialysis hour | P | per 24-h remoteness from 1<sup>st</sup> dialysis hour | P |
| HR, bpm | -0.76(-0.95 to -0.57) | <0.0001 | +0.02(-0.05 to 0.11) | 0.495 | +0.57(0.26-0.87) | <0.0001 | +0.20(-0.17 to 0.57) | 0.285 |
| rMSSD, ms | +0.20(0.12-0.29) | <0.0001 | -0.05(-0.09 to -0.001) | 0.044 | -1.41(-1.67 to -1.15) | <0.0001 | +0.10(-0.41 to 0.60) | 0.710 |
| HF power, s<sup>2</sup> | +0.04(-0.002 to 0.09) | 0.061 | -0.006(-0.03 to 0.02) | 0.588 | -0.38(-0.50 to -0.25) | <0.0001 | +0.24(0.06-0.42) | 0.010 |
| LF power, s<sup>2</sup> | +0.02(-0.002 to 0.05) | 0.073 | +0.002(-0.010 to 0.02) | 0.703 | -0.02(-0.08 to 0.05) | 0.639 | -0.04(-0.04 to 0.12) | 0.322 |
| LF/HF | -0.004(-0.01 to 0.003) | 0.246 | -0.001(-0.005 to 0.003) | 0.504 | +0.04(0.01 to 0.06) | 0.002 | -0.01(-0.03 to 0.01) | 0.157 |
| SD<sub>1</sub>, ms | +0.14(0.08-0.21) | <0.0001 | -0.03(-0.07 to -0.001) | 0.043 | -1.00(-1.19 to -0.81) | <0.0001 | +0.07(-0.29 to 0.42) | 0.709 |
| SD<sub>2</sub>, ms | -0.04(-0.26 to 0.18) | 0.706 | +0.27(0.17-0.37) | <0.0001 | +0.87(0.37-1.37) | 0.001 | -0.35(-0.81 to 0.12) | 0.148 |
| SD<sub>12</sub> | -0.003(-0.008 to 0.002) | 0.219 | -0.0006(-0.003 to 0.002) | 0.597 | -0.06(-0.07 to -0.05) | <0.0001 | +0.02(0.004-0.04) | 0.017 |
| SamEn | +0.006(-0.005 to 0.018) | 0.287 | +0.01(0.005 to 0.02) | <0.0001 | +0.007(-0.022 to 0.037) | 0.624 | -0.005(-0.047 to 0.036) | 0.795 |
| RenEn | +0.002(-0.002 to 0.005) | 0.345 | +0.004(-0.0005 to 0.008) | 0.088 | +0.02(0.002 to 0.04) | 0.030 | -0.01(-0.05 to 0.03) | 0.764 |

Sensitivity analysis, after exclusion of 3-minute epochs that started in the second half of an hour and thus violated equal intervals assumption provided consistently similar results (Table 8).

**Table 8.** Sensitivity analysis after exclusion of epochs starting in the 2^nd^ half of an hour: Association of the dialytic cycle with HRV.

| HR / HRV<br>metric | Dialysis & postdialysis (phases 1-2) |  | Regular interdialytic phases 3-4 |  | 2nd day interdialytic extension<br>(phases 5-6) |  | Nonadherent interdialytic extension<br>(phases 7-8) |  |
| --- | --- | --- | --- | --- | --- | --- | --- | --- |
|  | per 24-h remoteness<br>from 1<sup>st</sup> dialysis hour | P | per 24-h remoteness from<br>1<sup>st</sup> dialysis hour | P | per 24-h remoteness<br>from 1st dialysis hour | P | per 24-h remoteness<br>from 1st dialysis hour | P |
| HR, bpm | -0.60(-0.79 to -0.41) | <0.0001 | +0.02(-0.06 to 0.10) | 0.612 | +1.1(0.63-1.51) | <0.0001 | -0.06(-0.55 to 0.45) | 0.828 |
| rMSSD, ms | +0.20(0.10 to 0.30) | <0.0001 | -0.07(-0.11 to -0.02) | 0.006 | -1.51(-1.80 to -1.23) | <0.0001 | +0.33(-0.24 to 0.91) | 0.257 |
| HF power, s<sup>2</sup> | +0.04(-0.008 to 0.09) | 0.103 | -0.01(-0.03 to 0.01) | 0.332 | -0.36(-0.50 to -0.23) | <0.0001 | +0.26(0.06-0.47) | 0.013 |
| LF power, s<sup>2</sup> | +0.02(-0.008 – 0.05) | 0.176 | +0.003(-0.010-0.02) | 0.665 | +0.04(-0.04 to 0.12) | 0.319 | +0.07(-0.12 to 0.15) | 0.093 |
| LF/HF | -0.006(-0.01 to 0.003) | 0.180 | -0.002(-0.006 to 0.002) | 0.349 | +0.05(0.02 to 0.08) | 0.001 | -0.01(-0.04 to 0.01) | 0.377 |
| SD<sub>1</sub>, ms | +0.14(0.07-0.21) | <0.0001 | -0.05(-0.08 to -0.01) | 0.005 | -1.07(-1.28 to -0.87) | <0.0001 | +0.23(-0.17 to 0.65) | 0.257 |
| SD<sub>2</sub>, ms | -0.20(-0.43 to 0.04) | 0.103 | +0.29(0.18-0.39) | <0.0001 | +0.52(-0.13-1.18) | 0.119 | -0.87(-1.62 to -0.12) | 0.023 |
| SD<sub>12</sub> | -0.004(-0.009-0.0009) | 0.111 | -0.001(-0.004-0.001) | 0.275 | -0.06(-0.08 to -0.05) | <0.0001 | +0.02(0.004-0.04) | 0.017 |
| SamEn | +0.003(-0.009-0.02) | 0.595 | +0.010(0.004-0.015) | 0.001 | 0.006(-0.028 to 0.040) | 0.733 | +0.01(-0.033 to 0.055) | 0.631 |
| RenEn | +0.002(-0.002-0.006) | 0.243 | +0.004(-0.0003-0.008) | 0.069 | +0.02(0.002 to 0.04) | 0.030 | -0.01(-0.05 to 0.03) | 0.764 |

## Discussion

This continuous 14-day ECG monitoring study of incident dialysis patients revealed several important findings. First, we observed a significant association of dialytic phase with cardiac arrhythmias, with VT being more frequent during dialysis or within 6 hours post-dialysis. In the time-series analysis adjusted for cardiovascular disease and major risk factors, we demonstrated an independent association of incident VT events with significantly increased heart rate, confirming the importance of sympathetic activation as a VT trigger. Secondly, we found that the dialysis schedule dramatically influenced the autonomic tone. Regular dialytic schedule (every other day dialysis) preserved physiological circadian rhythm in heart rate and HRV, whereas the extension of the interdialytic period for the second day abolished circadian rhythm and displayed a steady deterioration: progressively decreasing parasympathetic and increasing sympathetic tone. Finally, we showed that associations of clinical and demographic characteristics with heart rate and HRV time series are different in different phases of the dialytic cycle, highlighting difficulties in assessment of autonomic nervous system status in ESKD patients on dialysis.

### Dialysis phase and incident cardiac arrhythmias during ECG monitoring

An association between the hemodialysis schedule and SCD rate is well-known: SCD is more frequent after the long interdialytic interval, during dialysis, or within 6-12 hours after the end of a hemodialysis session^6-8^. Our observation was consistent with implantable loop recorder (ILR) studies^33^ that confirmed the same pattern: VT events were more frequent during and post-dialysis. This finding supports the hypothesis of an interaction between the pre-existing electrophysiological substrate and plastic chemical exposure during dialysis resulting in increased risk of life-threatening VT^10^. Accessibility to external defibrillators during and within 6-12 hours after dialysis can be potentially life-saving. A randomized clinical trial is needed to test this hypothesis.

Importantly, after rigorous adjustment for cardiovascular disease and its risk factors in comprehensive ARCH/GARCH models, we observed an association of increased heart rate with paroxysmal VT events. Unfortunately, the patient’s activity at the times of increased heart rate and VT events was not known. The sympathetic drive is a well-known trigger of VT and SCD^12^. Administration of beta-blockers^34^ can be potential preventive intervention in such patients.^3^

### Dynamic changes in autonomic tone in the dialytic cycle

Despite multiple studies reporting unfavorable effects of the long 2-day interdialytic “weekend,” ^6-8^ hemodialysis in the US is typically prescribed three times per week, with two 1-day and one 2-day interdialytic intervals between treatment sessions, because dialysis centers in the US are closed on Sundays. Our study is the first long-term study of heart rate and HRV time-series across multiple dialytic cycles, allowing direct paired comparison of the effect of the second-day interdialytic interval extension. Our findings added to the strong mounting evidence of the harmful consequences of the long interdialytic interval. While routine dialytic schedule (on every other day) preserved relatively normal cardiovascular autonomic control, the second day without dialysis was characterized by parasympathetic withdrawal and steady increase in sympathetic predominance, which may explain the previously observed increased rate of SCD after long interdialytic interval.^6-8^ Mounting evidence of a harmful effect of the two-day interval without dialysis suggests that everyday^35^ or every other day dialysis^36^ should be considered as a preferred treatment schedule.

Having long interdialytic weekend was especially disadvantageous to women, blacks, participants with the history of CVD and a large number of comorbidities (high CCI). Of note, in fully adjusted ARCH/GARCH models, subjective feeling of a long recovery after dialysis was independently associated with greater autonomic disbalance, suggesting that a simple question “How long does it take you to recover from a dialysis session?” can carry meaningful information, useful for treatment management.

### Effect of missing dialysis

A surprisingly large subgroup of our study participants missed 1-2 dialysis sessions during ECG monitoring, providing a unique opportunity to observe the effect of an ultra-long interdialytic interval. As expected, heart rate further slowed, HRV progressively diminished, and circadian rhythm in HRV remained vanished or distorted, due to deepened autonomic disbalance. Of note, patients who missed dialysis were characterized by prominent bradycardia. Several recent studies reported bradyarrhythmias during monitoring,^5, 9, 33, 37^ but did not comment (or did not have available data) on patients’ adherence to treatment. Improvement of patient adherence, avoiding missing dialysis sessions can improve patients’ outcomes.

### Heart rate variability in ESKD patients on dialysis

HRV reflects dynamic bidirectional interaction between the heart and the respiratory system, regulated by the autonomic nervous system^14^, estimating sympathovagal balance.^38^ Our study highlights the unique features of HRV-manifestation of autonomic imbalance in ESKD patients presenting with bradycardia due to direct negative chronotropic and dromotropic effects of hyperkalaemia, hypocalcaemia, and uremic toxins. Autonomic imbalance can diminish the cholinergic anti-inflammatory pathway and lead to an exaggerated cytokine response, promoting inflammation, which is unfavorable for ESKD patients^39^. As others,^40^ we observed heart rate increase during and immediately after dialysis.^41^ In our study participants, autonomic imbalance manifested largely by parasympathetic withdrawal (decrease in rMSSD, HF power, and SD_1_). Unlike in CHF studies,^42, 43^ we observed LF/HF ratio less than 1, likely because of impaired baroreflex sensitivity, or blunted sympathetic response in our study participants. Moak *et al*.^13^ showed that LF power reflects baroreflex-mediated changes in cardiovagal and sympathetic noradrenergic outflows. In the case of baroreflex failure, LF power is reduced, regardless of the status of cardiac sympathetic innervation. Normally, acute removal of fluid during dialysis session activates the baroreflex. Several groups of investigators^44-47^ observed worse clinical outcomes in ESKD patients with the blunted sympathetic response and low (less than 1) LF/HF power ratio, which was characteristic of our patient population. Blunted sympathetic response on volume removal can be genetically determined by the polymorphism in the gene for angiotensin-converting enzyme.^40^ Inadequate baroreflex sensitivity can cause intradialytic hypotension,^48^ a well-known risk marker of adverse clinical outcomes in ESKD.

### Circadian rhythm in heart rate and HRV

Our finding of disrupted circadian rhythm is in agreement with previously reported autonomic dysregulation during normal wake/sleep cycle in ESKD.^49^ Two mechanisms^50^ are responsible for circadian rhythm in heart rate: (1) central circadian clock in the suprachiasmatic nucleus in the hypothalamus, acting via the autonomic nervous system, and (2) a local circadian clock in the heart itself. In our study participants, the circadian rhythm in heart rate remained relatively preserved during the prolonged interdialytic interval, whereas the circadian rhythm in HRV was largely abolished. Changes in the acrophase of circadian rhythm in heart rate during ultra-long interdialytic (missed dialysis) interval (peak in the afternoon instead of morning hours) may reflect switch of the circadian clock from central to local regulatory mechanisms. A better understanding of circadian rhythms in heart rate and HRV may help to improve medical management and clinical outcomes in ESKD patients on dialysis. Chan *et. al*.^35^ studied the effect of daily dialysis versus every-other-day dialysis and found that daily dialysis improved vagal modulation of the heart and increased short-term HRV, which is consistent with our study findings. Further studies of the interplay between circadian rhythm and dialytic cycle are needed, to develop optimal treatment schedule for ESKD patients.

### Time-series analysis and periodic regression

In this study, we were the first to appropriately model associations of clinical characteristics, incident arrhythmias, circadian and dialytic cycles with heart rate and HRV time series using robust ARCH/GARCH models. ARCH models allow modeling of the time-dependent mean and time-dependent variance of studied outcomes (heart rate / HRV time series), which accurately reflect volatility that can arise in response to the dialysis procedure or other unmeasured factors during long-term ECG monitoring (various types of activity). ARCH methods are currently successfully applied in the statistical finance field, to model the volatile stock market. Meaningful results of our study call for broader use of ARCH models in the analysis of physiological data time-series.

Importantly, we used explicit (sin’X, cos’X) periodic regression, but not a so-called “cosinor regression,” which may drastically overestimate significance due to inappropriate correction for the degree of freedom. Use of explicit periodic regression allowed us to fit both circadian and dialytic cycles together and observe a longitudinal trend in HRV after removal of the circadian pattern.

### Strengths and Limitations

The strengths of this study arise from long-term continuous ECG monitoring, providing a unique opportunity for our pioneering study of multiple 24-hour and dialytic cycles for the same participants, significantly improving robustness and reducing error in estimations. Other strengths include the use of advanced statistical modeling of a time-series (ARCH/GARCH) and appropriate use of explicit periodic regression.

However, limitations of the study should be considered. Use of a single-lead ECG poses objective challenges for discrimination of supraventricular arrhythmia from normal sinus rhythm. Everyday physical activity manifests by noise and artifacts. To ensure analysis of normal sinus rhythm, we implemented vigorous quality control procedures, which included semi-automated analysis of the data, and manual review of ECGs. To improve the quality of included ECG data, we elected to study 3-minute epochs of continuously normal (uninterrupted) clean sinus rhythm.

Impaired baroreflex sensitivity can be measured by heart rate turbulence (HRT), which was not measured in this study. Further study of HRT in ESKD patients on dialysis is needed. Results of frequency-domain HRV should be considered with caution. We measured HRV on each 3-minute segment, which at least partially explains the differences between HF and LF power reported in our study as compared with the 24-hour power spectrum analyses.^42, 43^ To overcome limitations of isolated HRV metrics, we evaluated several HRV measures presumably reflecting parasympathetic tone (rMSSD, SD_1_, HF power), and sympathovagal balance (SD_2_, SD_12_, LF/HF ratio). Consistent findings across the full set of HRV metrics increases the validity of our results.

The size of this study is relatively small. Nevertheless, the study was sufficiently powered for the outcome: time-series of heart rate and HRV. On average, the study participant had 156±55 hours of data. Statistical power of the periodic regression and time-series analysis was above 0.8.

The assumption of equal intervals between 3-minute epochs was violated for approximately 10% of epochs. To address this limitation, sensitivity analyses were performed and epochs that started in the second half of an hour were excluded from ARCH models, which did not change the association of dialytic cycle with HRV time-series (Table 8). Only 4 study participants missed dialysis, and therefore, results of this subgroup analyses should be considered with caution. The reason for the missed dialysis sessions was unknown. It is possible that participants missed dialysis because of travel, which may explain the shift in circadian rhythm acrophase.

### Funding Sources

The PACE Study was supported by NIDDK grant R01DK072367 (to R.S.P.). This study was supported in part by the NHLBI grant HL118277 (to L.G.T.).

### Disclosures

None

